# Plant growth promoting activity of bacteria isolated from Asian rice (*Oryza sativa* L.) are plant subspecies dependent

**DOI:** 10.1101/2021.12.21.473765

**Authors:** Nasim Maghboli Balasjin, James S. Maki, Michael R. Schläppi, Christopher W. Marshall

## Abstract

Asian rice is one of the most important crops because it is a staple food for almost half of the world’s population. Rice has two subspecies, *JAPONICA* and *INDICA*. To have production of rice keep pace with a growing world population, it is anticipated that the use of fertilizers will also need to increase, which may cause environmental damage through runoff impacts. An alternative strategy to increase crop yield is the use of plant growth promoting bacteria. Thousands of microbial species can exist in association with plant roots and shoots, and some are critical to the plant’s survival. We isolated 140 bacteria from rice and investigated whether *JAPONICA* and *INDICA* rice subspecies were positively influenced by these isolates. The bacterial isolates were screened for their ability to solubilize phosphate, a known plant growth promoting characteristic, and 25 isolates were selected for further analysis. These 25 phosphate solubilizing isolates were also able to produce other potentially growth-promoting factors. Five of the most promising bacterial isolates were chosen for whole genome sequencing. Four of these bacteria, isolates related to *Pseudomonas mosselii, Microvirga* sp., *Paenibacillus rigui* and *Paenibacillus graminis*, improved root and shoot growth, root to shoot ratio, and increased root dry weights of *JAPONICA* plants but had no effect on growth and development of *INDICA* plants. This indicates that while bacteria have several known plant growth promoting functions, their effects on growth parameters can be plant subspecies dependent and suggest close relationships between plants and their microbial partners.

## Introduction

Plant-microbe interactions are divided into three groups including pathogenic, symbiotic, and associative. Each of these interactions can affect plant physiology such as nutrition level, growth and development, and defense mechanisms (1). These interactions are beneficial to both plants and microorganisms (2). Plant-associated bacteria that benefit development of plants through direct and indirect mechanisms are known as plant growth promoting bacteria (PGPB) (3). Direct mechanisms of plant growth promotion include phosphorus and zinc solubilization, nitrogen fixation, plant hormone (phytohormone) production, and indirect mechanisms include antifungal activity, lytic enzyme production, siderophores and ammonia production (3).

There are a variety of PGPB that inhabit the area near plant roots (rhizosphere), the root surface (rhizoplane), and inside the root (endosphere) (4). PGPB that live inside plant tissues are known as endophytes (5). Plants can attract specific bacteria from the soil environment to live on or inside their roots, to benefit the plants by providing and solubilizing nutrients indicating the importance of the rhizoplane and the endosphere bacterial communities (6). On the other hand, plant roots provide carbon metabolites and nitrogen to bacteria (2). The most common plant growth promoting bacteria belong to the genera *Bacillus, Pseudomonas, Enterobacter, Acinetobacter, Burkholderia, Arthrobacter*, and *Paenibacillus*. These bacteria are known to enhance plant immunity aginst pathogens and provide phytohormones, soluble phosphate, and/or nitrogen (7, 8, 9).

Asian rice (*Oryza sativa* L.) is one of the main staple foods for almost half of the world’s population (10). It is estimated that by 2050 food demand will be heightened due to an increasing global population (11). Therefore, the rate of fertilizer use will increase, causing environmental problems (12). One promising solution is to create environmentally friendly bio-fertilizers containing plant growth promoting bacteria. Phosphorus is the second most important nutrient, after nitrogen, for plants but is typically insoluble in the soil (13).For plant roots to take up phosphate, insoluble inorganic phosphate needs to be converted to soluble phosphate (13, 14). Phosphate solubilizing bacteria (PSB) play an important role because they secrete gluconic and keto-gluconic acids and phosphatases that release soluble phosphates into the soil (14, 15, 16). It has previously been shown that *Anabaena*, *Azospirillum, Rhodobacter* and *Streptomyces* species can promote *O. sativa* growth, but further characterization is needed to expand the known species capable of growth promotion, to understand their mechanisms of action, and their rice specificity for rice subspecies (4, 15, 17). Therefore, discovering and characterizing phosphate solubilizing bacteria can improve crop productivity with less environmental impact than traditional fertilizers.

In this study, we screened for plant growth promoting bacteria by isolating bacteria from the surface and inner parts of leaves and roots of *INDICA* and *JAPONICA* rice, the two subspecies of *O. sativa*. Bacterial isolates were evaluated for plant growth promoting phenotypes and the genomes of five PGPB were sequenced. Finally, we separately evaluated the effect of the sequenced isolates on growth and development of *JAPONICA* and *INDICA* rice accessions. To the best of our knowledge, this is the first work evaluating the influence of *O. sativa* endophytes, that are isolated from both *INDICA* and *JAPONICA*, on the growth and development of each of the two subspecies. This study investigated the influence of endophytes isolated from one rice subspecies (*JAPONICA*) on the other subspecies (*INDICA*), and *vice versa*.

## Materials and methods

### Plant growth and bacterial isolation from roots and leaves

We used accessions from the two subspecies of *O. sativa*: Krasnodarskij 3352, (USDA # PI311787) from the *temperate japonica* subpopulation representing the *JAPONICA* subspecies; and Carolino 164 (USDA # PI311654) from the *aus* subpopulation representing the *INDICA* subspecies. Following the methods of (4), *INDICA* and *JAPONICA* seeds were surface sterilized with minor modifications. Briefly, seeds were dehulled and immersed in ethanol (70% v/v in dH_2_O) for 1 min, rinsed with sterile water (3X), immersed in sodium hypochlorite (70% v/v in dH_2_O) for 5 min and rinsed again with sterile water (3X). After drying surface sterilized seeds on autoclaved filter papers, seeds were put on sterile agar-solidified Murashige-Skoog (MS) medium (4) for germination and incubated at 37°C (2 d). Germinated seeds were transferred to a growth chamber with cycles of 12 h of light (28°C) and 12 h of dark (24°C) and after 10 d of growth, seedlings were transplanted into soil pots.

Isolation of bacteria from the roots (rhizoplane and endosphere) and leaves (phyllosphere) during vegetative growth (when plants were two weeks old) was done as described previously (18). Roots and leaves of both rice subspecies were rinsed with a sterile phosphate-buffered saline (PBS; 8 g NaCl, 0.2 g KCl, 1.44 g Na_2_HPO_4_, 0.24 g KH_2_PO_4_, pH = 7.4 with HCl, total volume = 1 L) to remove the attached soil from roots and other contaminants from leaves. Rinsed roots and leaves were sonicated 3X in sterile PBS and surface sterilized following the methods of (19). Briefly, the roots and leaves were rinsed with sterile water, then soaked in ethanol (75% v/v in dH_2_O) for 1 min and again rinsed with sterile water. Next, they were soaked in sodium hypochlorite (1% v/v in dH_2_O) for 1 min and finally washed-with sterile water. Sterile roots and leaves were crushed in PBS with a sterile mortar and pestle. The PBS that contained crushed roots and leaves was transferred into 1.5 mL sterile tubes and centrifuged (12,074 x g, 10 min). 100 μL of the supernatant was plated on nutrient agar (NA) media (Difco) and incubated at 30°C. Cultures were purified by repeatedly streaking bacteria onto NA and incubating them at 30°C until only one colony type remained.

### Colony and cell morphology of isolated bacteria from roots and leaves

Colony and cell morphology of isolated bacteria were determined through observing purified cultures and gram staining, respectively. Colony morphology characteristics include form, elevation, size, color, and margin were investigated (https://laboratoryinfo.com/colony-morphology-of-bacteria/; last accessed during the Spring of 2019).

Pure cultures (OD_600_ = 0.6) used for agar assay methods and spot inoculations were grown in nutrient broth (NB) medium at 30°C for 24 h.

### Mineral phosphate solubilization activity test

Pure cultures were grown in nutrient broth (NB) media at 30°C for 24 h. An agar assay method was used to screen the phosphate solubilizing ability of bacterial isolates. Following the methods of (20), the National Botanical Research Institute’s phosphate (NBRIP) growth medium containing glucose (10 g L^-1^), Ca_3_(PO_4_)_2_ (5 g L^-1^), MgCl_2_.6H2O (5 g L^-1^), MgSO_4_.7H_2_O (0.25 g L^-1^), KCl (0.2 g L^-1^) and (NH_4_)_2_SO_4_ (0.1 g L^-1^) was used for this approach. Two μL of each isolated bacterium (OD_600_ = 0.6) was spot inoculated onto NBRIP (National Botanical Research Institute’s Phosphate) medium in three replicates. Plates were incubated at room temperature for 14 d. After two weeks plates were examined for phosphate solubilization activity as shown by a clear halo around the colony. The phosphate solubilization index (20) was calculated from three independent experiments as follows:

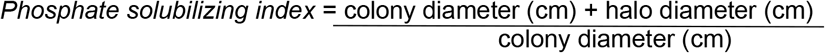

### Screening phosphate solubilizing bacteria (PSB) for other plant growth promoting features

Both direct and indirect mechanisms of plant growth promoting screening tests were done on PSB using the following culture-based assays (also listed in Supplementary Table S1):

### Zinc solubilization

Tris minimal medium (Tris-HCl 6.06 g L^-1^, NaCl 4.68 g L^-1^, KCl 1.49 g L^-1^, NH_4_Cl 1.07 g L^-1^, Na2SO_4_ 0.43 g L^-1^, MgCl_2_.2H_2_O 0.2 g L^-1^, CaCl_2_.2H_2_O 30 mg L^-1^, agar 15 g L^-1^) with 0.1% (w/v in distilled water) zinc sulfate (ZnSO_4_) was prepared and 2 μL of each isolated strain (OD_600_= 0.6) in PBS was spot inoculated onto the mentioned medium in three replicates. Plates were incubated at 30°C for 14 d. After two weeks plates were examined for zinc solubilization activity. Similar to phosphate solubilization, a halo zone around each bacterium was confirmation for zinc solubilization (21). This experiment was repeated twice, independently.

### Nitrogen fixation

For confirmation of nitrogen fixation, through culture-based assays, 2 μL of PSB (OD_600_ = 0.6) were spot inoculated onto nitrogen-rich yeast extract-mannitol agar containing Congo Red dye (CR-YMA) and incubated at 30°C for 5 d. Weak-absorption of Congo Red dye by colonies was confirmation for nitrogen fixation. Bacterial colonies with colors ranging from white to pale white-pink were able to fix nitrogen (22) due to cleavage of the azo bond (-N=N-) in Congo Red (23). This experiment was repeated three times, independently. *Rhizobium* sp. and *Staphylococcus aureus* were used as positive and negative controls, respectively.

### Indoleacetic acid (IAA) production

A colorimetric assay was done using the ferric chloride-perchloric acid reagent (FeCl_3_-HClO_4_) to detect IAA production. Indole compounds were quantified in precursor L-tryptophan medium. Nutrient Broth-M26 (NaCl 5 g L^-1^, peptone 10 g L^-1^, and beef extract 10 g L^-1^) was used to grow bacteria for 24 h on a shaker (150 rpm) at 28°C. 100 μL of the cultures were inoculated into 10 mL of liquid minimal salt (MS) medium (KH_2_PO_4_ 1.36 g/l, Na_2_HPO_4_ 2.13 g L^-1^, MgSO_4_.7H_2_O 0.2 g L^-1^, and trace elements (containing citric acid 5 g/100 mL, ZnSO_4_.7H_2_O 5 g/100 mL, FeSO_4_.7H_2_O 4.75 g/100 mL, Fe(NH_4_)2(SO_4_)2.6H_2_O 1 g/100 mL, CuSO_4_.5H_2_O 250 mg/100 mL, MnSO_4_.H_2_O 50 mg/ 100 mL, H_3_BO_3_ 50 mg/ 100 mL and Na_2_MoO_4_.2H_2_O 50 mg/ 100 mL in 100 mL distilled water) 1 mL) supplemented with 5 mM L-tryptophan and incubated at 28°C for 48 h on a shaker (150 rpm). L-tryptophan medium contained glucose (10 g L^-1^), L-tryptophan (1 g L^-1^), and yeast extract (0.1 g L^-1^) in 100 mL water and was filter sterilized through a 0.2 μm membrane (Whatman Syringe Filters). After 48 h of growth, 1.5 mL of bacterial solution were centrifuged at 8,870 x g for 5 min in a microfuge. One mL of the supernatant was mixed with 2 mL FeCl_3_-HClO_4_ reagent and after 25 min, the mixture’s optical density (OD) was measured using a UV-spectrophotometer at 530 nm. A standard curve (with concentrations of 0-300 μg/mL) was created for calculating the microgram of IAA per mL of the mixture (24). This experiment was repeated twice, independently.

### Gibberellic acid (GA) production

GAs are other important phytohormomes regulating plant growth, seed germination, and stem elongation (25). Following the methods of (25) and (26), the GA production detection assay was done with minor modifications. Briefly, PSB were freshly grown in NB (Difco) medium for 24 h at 30°C, then incubated at room temperature for one week. After growth, 1.5 mL of bacterial suspensions were centrifuged (3,942 x g) for 10 min in a microfuge. One mL of supernatant was transferred into 15 mL tubes and 2 mL of zinc acetate solution (zinc acetate 21.9 g L^-1^, glacial acetic acid 1 mL, distilled water up to 100 mL) was added to the supernatant. Two mL of potassium ferrocyanide solution (10.6% w/v in distilled water) was added to the 15 mL tubes containing supernatant and zinc acetate solution. Tubes were centrifuged (7,168 x g, 10 min) and 1 mL of supernatant was transferred to another 15 mL tube and 5 mL HCl solution (30% v/v in distilled water) was added. Tubes were incubated 75 min at 28-30°C after which OD_254_ measurements were taken. A standard curve (with concentrations of 0-1000 μg mL^-1^ of GA) was created for calculating the microgram of gibberellic acid per milliliter of the mixture (25, 26). This experiment was repeated twice, independently.

### 1-aminocyclopropane-1-carboxylic acid (ACC) deaminase activity

A culture-based assay was done for ACC deaminase activity as described previously (21). Bacterial isolates were spot inoculated in three replicates and grown on Dworkin & Foster (DF) minimal salt medium (DF salts per liter: 4.0 g KH_2_PO_4_, 6.0 g Na_2_HPO_4_, 0.2 g MgSO_4_.7H_2_O, 2.0 g glucose, 2.0 g gluconic acid and 2.0 g citric acid with trace elements: 1 mg FeSO_4_.7H_2_O, 10 mg H_3_BO_3_, 11.19 mg MnSO_4_.H_2_O, 124.6 mg ZnSO_4_.7H_2_O, 78.22 mg CuSO_4_.5H_2_O, 10 mg MoO_3_, pH 7.2) supplemented with 3 mM ACC as the sole nitrogen source. Isolates that grew on the plates were able to produce ACC deaminase (21). This experiment was repeated three times, independently.

### Antifungal activity

A culture-based assay against the fungal pathogen *Magnaporthe grisea* was done as described previously (27), with minor modifications. Briefly, each isolate was cultured in two parallel lines (2-2.5 cm apart from each other) on potato dextrose agar (PDA) medium. *Magnaporthe grisea* causing rice blast disease was isolated from leaves of rice plants grown in rooftop paddies outside of the Schläppi lab and was spot cultured in the center of PDA plates (between two parallel bacterial lines, 0.5-1 cm away from either sides). Control plates only had the fungus without bacteria. Plates were incubated at 28°C for five days. The radial growth of the fungus was examined to observe growth inhibition by each bacterium. Percentage of growth inhibition from three independent experiments was calculated and compared with the radial growth of fungus in control plates using the formula shown below with the following parameters: I = inhibition percentage; C = radial growth in control plate; T = radial growth in plates with bacterial isolates (27):

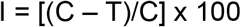

### Lipase production

An agar assay method was used to detect bacteria with lipase production ability. Luria-Bertani (LB) medium (tryptone 10 g, NaCl 10 g, yeast extract 5 g, agar 15 g) was prepared in 490 mL distilled water and supplemented with 1% tween-20. Two μL of each isolated strain was spot inoculated on the mentioned medium in duplicate. One set of plates was incubated at 28°C for 3-4 d. Lipase production was detected by observing precipitation around colonies (28). This experiment was repeated three times, independently.

### Casein and gelatin hydrolyzing

Protein hydrolysis was tested using both casein and gelatin media. Casein agar medium contained NB (Difco) with 1.5% (w/v) agar. 50 mL of evaporated milk with 50 mL of sterile water were added to the sterile NB with agar (final volume = 1000 mL) and 2 μL of each isolated strain (OD_600_ = 0.6) was spot inoculated onto the medium in duplicate. Plates were incubated at 28°C for 24 h. Casein hydrolysis was examined by observing a halo around colonies (29). Gelatin medium contained 3 g L^-1^ beef extract, 5 g L^-1^ peptone and 60 g L^-1^ gelatin in 1 L of distilled water. Bacterial solutions were stabbed into test tubes containing gelatin with inoculating needles. Tubes were incubated at 28°C for two weeks. Bacteria with gelatinase activity were able to liquify the medium (30). This experiment was repeated three times, independently.

### Siderophore production

An agar assay method was used to detect bacteria producing siderophores that might help plants with iron uptake from the soil (30), as described previously (31). Specifically, chrome azurol S (CAS) agar was prepared by mixing four sterile solutions. Solution 1, the Fe-CAS indicator solution, contained 10 mL of 1 mM FeCl_3_.6H_2_O (in 10 mM HCl), 50 mL of an aqueous solution of CAS (1.21 mg mL^-1^), and 40 mL of an aqueous solution of hexadecyl-trimethylammonium bromide (HDTMA) (1.82 mg mL^-1^). Solution 2, the buffer solution, contained 30.24 g piperazine-N,N’-bis[2-ethanesulfonic acid] (PIPES) dissolved in 750 mL of salt solution (100 mL of MM9 salt solution dissolved in 750 mL of distilled water). 50 mL water was added to bring the final volume to 800 mL and 1.5% (w/v) agar was added. To dissolve PIPES in the salt solution, 50% KOH was added until the pH of the solution was 6.8. Solution 3 contained 2 g glucose, 2 g mannitol, and trace elements (same ingredients mentioned in IAA production method section) which were dissolved in 70 mL distilled water. Solution 4 contained 10% (w/v) casamino acids, which was filter sterilized (0.2 μm membrane, (Whatman Syringe Filters)). All solutions, except solution 4, were autoclaved. After cooling to 50°C, solutions 3 and 4 were mixed, then the buffer solution (solution 2), was added. The combined solutions were added to solution 1. The prepared medium (blue to dark green color) was poured into plates and 2 μL of each isolated strain (OD_600_ = 0.6) was spot inoculated onto the medium in duplicates. Plates were incubated at 30°C for 24 h. Siderophore production was examined by observing an orange halo around colonies (31, 32). This experiment was repeated three times, independently.

### Cellulase production

The medium used for this test was the same as for lipase production, except that instead of tween-20, the medium was supplemented with 1% carboxymethyl cellulose (CMC). Two μL of each isolated strain (OD_600_ = 0.6) (*E. coli* = negative control) was spot inoculated onto the medium in duplicate. Plates were incubated at 28°C for 3-4 d. Cellulase production was detected through staining and de-staining with 0.1% Congo Red (w/v in distilled water) for 15 min and 1M NaCl for 15 min, respectively (33). This experiment was repeated three times, independently.

### Ammonia (NH_3_) production

A colorimetric assay was done following the methods of (33) with minor modifications. Freshly grown bacterial isolates were inoculated in 10 mL peptone water (15 g peptone water in 1000 mL distilled water) and incubated at 28°C on a shaker (200 rpm) for 48 h. After incubation, 1 mL of each bacterial sample was transferred to a 1.5 mL tube and centrifuged for 10 minutes (2,218 x g) in a microfuge. The supernatant was transferred to another 1.5 mL tube and 100 μL Nessler’s reagent [10% HgI2; 7% KI; 50% aqueous solution of NaOH (32%)] was added to each tube and incubated at room temperature for 30 min. Samples with the ability of ammonia production turned a yellow to orange color. Optical density of each sample was measured in a UV-spectrophotometer at 520 nm. A standard curve (with concentrations of 0-300 μg mL^-1^ using ammonia carbonate) was created for calculating the μg of NH_3_ per mL of the mixture solution (33). This experiment was repeated three times, independently.

### *In vitro* assay for salt tolerance

Bacterial isolates were spot inoculated in three replicates on Luria-Bertani (LB) medium (tryptone 10 g L^-1^, NaCl 10 g L^-1^, yeast extract 5 g L^-1^, agar 15 g L^-1^) supplemented with 0%, 2%, 4%, 6% and 8% NaCl. LB plates were incubated at 28°C for 5 d. Plates were examined for bacterial growth on LB with different NaCl concentration (21). This experiment was repeated three times, independently.

### Whole genome sequencing of 5 candidate PSB

Five PSB isolates from the *INDICA* endosphere, *INDICA* phyllosphere, and *JAPONICA* phyllosphere were chosen based on differing colony morphologies and their ability to produce plant hormones for whole genome sequencing. Nucleic acid extraction of the 5 PSB was done using the Qiagen DNeasy Blood and Tissue Extraction kit according to the manufacturer’s instructions. The DNA concentration of each sample was measured with a Nanodrop spectrophotometer. The PSB samples were then sent to the Microbial Genome Sequencing center (www.migscenter.com) for whole genome sequencing. Libraries were prepared as described previously (34) and sequenced on an Illumina NextSeq 550 yielding 151-bp paired end reads.

### Sequence analysis and annotation of sequenced PSB

Bacterial sequences of 5 specific PSB were analyzed using KBase.us and data were deposited into the public narrative site https://narrative.kbase.us/narrative/52526. Sequence reads were quality checked with FastQC v0.11.9 (35).Genomes were assembled with SPAdes v3.13.0 (36) and annotated with RASTtk (37). Initial genome relatedness was determined by inserting a genome into a phylogenetic tree with FastTree2 (38) and calculating average nucleotide identity with FastANI (39) and JSpeciesW (40) on closest genome relatives. Taxonomic classification was done with the genome taxonomy database toolkit v1.6.0 (GTDB-tk) (41). A final tree was generated using the IQ-TREE2 v2.1.4 (42) implementation of GToTree v1.6.11 and its dependencies (43, 44, 45, 46, 47, 48).

### Phenotypic fingerprint of sequenced PSB

This test is specialized for characterizing bacteria phenotypically using the Biolog Gen III Microplate. The 5 sequenced PSB were characterized utilizing 71 carbon sources and 23 chemical sensitivity assays on 96 well microplates (Supplementary Fig. S1). Isolated bacteria were grown on NA medium and suspended in inoculating fluid (provided with the 96 well microplates) using sterile swabs. The turbidity of inoculating fluid (IF) was 95%. Then, 100 μL of the bacterial suspension were added to each well and microplates were incubated at 28°C for 2-3 d. Wells that were positive for a specific carbon source turned purple due to formazan (if absorbance determined in a microplate reader exceeded the control well) or were sensitive to a chemical material (if formazan absorbance was less than the control well) (www.biolog.com). This experiment was repeated three times.

### Antibacterial activity of the sequenced PSB against each other

Two culture based antibacterial activity assays, overlay-agar, and cross streak, were done to determine whether the 5 sequenced PSB interact with each other:

### (1) Overlay-Agar assay

Isolated bacteria were grown in nutrient broth (NB) media at 30°C for 24 h. Freshly grown bacteria (OD_600_ = 0.6), were spot inoculated on NA media and incubated at 30°C for 48 h. After 2 d of incubation, plates were inverted under the hood over filter papers soaked in chloroform for 15 min. Plates were put upright without lids to let extra chloroform evaporate. 10 mL of NB media (containing 1.5% agar at 50°C) mixed with 200 μL of an indicator strain (the organism to be tested for susceptibility) was poured on the surface of NBA plates and incubated at 30°C for 48-72 h. After incubation, inoculated bacteria were checked for zones of inhibition (49). This assay was repeated three times, independently.

### (2) Cross-streak agar assay

Isolated bacteria were grown in NB media at 30°C for 24 h. Freshly grown bacteria (OD_600_ = 0.6) were inoculated in a single 1 cm wide linear streak down the center of NA plates and this process was repeated for each bacterium. Plates were incubated at 30°C for 48 h. The organism to be tested for susceptibility, was streaked in lines perpendicular towards the bacterium that has been horizontally streaked (up to 1 mm). Plates were incubated at 30°C for 48-72 h. After incubation, plates were examined for zones of inhibition around the initial bacterium streak, and the width of each zone of inhibition was measured in mm (50). This assay was repeated three times, independently.

### Evaluating the influence of sequenced PSB on rice growth and development

The influence of five sequenced PSB on rice growth and development was evaluated by seed inoculation with bacterial suspensions [control plants: seeds inoculated in bacterial free KCl]. Shoot and root lengths were measured when plants were 8, 12 and 14 days old. Furthermore, when plants were two-week old, dry weight root/shoot ratio was determined to measure nutrient uptake ability (the higher the root/shoot ratio, the more nutrients are taken up by plants) (51).

### Shoot and root length

Bacterial cells were harvested by centrifugation and rinsed with sterile water, then resuspended in 0.85% KCl (biological saline) (52). *INDICA* and *JAPONICA* wild type seeds were surface sterilized as described earlier and soaked in the bacterial suspension (10^8^ cell mL^-1^) overnight (53). Soaked seeds in KCl solution were used as controls and counted as non-bacterial soaked seeds. Both soaked and non-bacterial soaked seeds were germinated in 1X MS media at 30°C for two days. Germinated seeds were transferred to autoclaved PCR strips placed in boxes filled with sterile water to grow hydroponically. The boxes were put in controlled chambers [12 h of light (28°C) and 12 h of dark cycle (24°C)] for 14 days). At day 10, water in boxes were replaced with ¼ Murashige and Skoog (MS) medium. Shoot and root lengths were measured when plants were 14 days old.

### Dry weight root/shoot ratio

Shoots and roots of 15 inoculated and uninoculated plants (two-week old plants) were isolated and put into a dry oven at 50°C temperature for 48 h (53). After two days of incubation, the dry weight root/shoot ratio of each of 15 plants per trial (total of 3 trials) were measured to evaluate the ability of plants to take up nutrients while they were inoculated with bacterium.

### Data availability

All sequence reads were deposited into the NCBI Sequence Read Archive (SRA) under BioProject PRJNA667792 and SRA accessions SRS7484588 - SRS7484592. All assemblies and annotations can be viewed and downloaded from the open access KBase narrative https://narrative.kbase.us/narrative/52526.

## Results

### Bacterial isolation and characterization

To identify plant growth promoting bacteria (PGPB) associated with rice plants, a total of 140 bacteria were isolated and purified from the phyllosphere, root endosphere, and root rhizoplane of two accessions representing the two *JAPONICA* and *INDICA* subspecies of rice (Supplementary Table S3). The colony morphology and Gram reaction of each isolated bacterium are listed in Supplementary Table S4. Because phosphate is the second most important macronutrient for plant growth (16), we reasoned that mineral phosphate solubilizing bacteria (PSB) associated with rice tissues might act as PGPB. Phosphate is mostly insoluble in the soil and therefore unavailable to plants (54). There are PSB in the soil with the ability of providing soluble phosphate through releasing organic acids and acid phosphatase (55). To identify PSB among the 140 isolates, bacteria were spotted onto agar plates containing insoluble phosphate (Ca_3_(PO_4_)_2_). Of the 140 isolates, 25 (18%) were able to solubilize mineral phosphate as shown by a halo around the bacterial colonies. The mean phosphate solubilization index was calculated for those 25 bacteria (Fig. 1 (A)). Isolate n00132 had the highest phosphate solubilization ability (mean phosphate solubilization index = 2.36 cm, *p* < 0.05; Supplementary Fig. S2 (C)).

**FIG 1.**
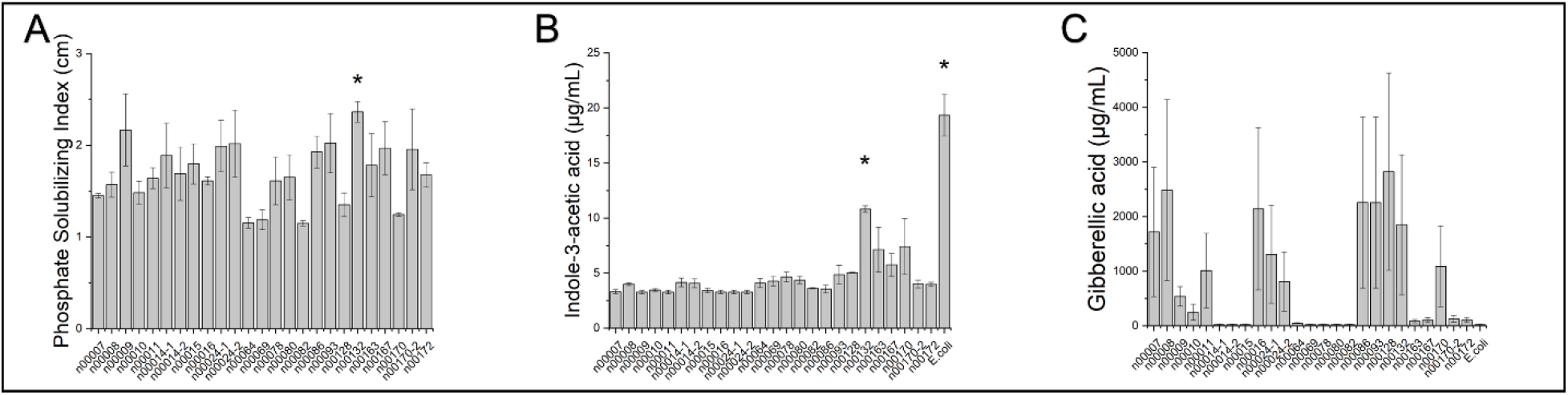
**(A)** Mean phosphate solubilization index of three trials. Phosphate solubilization activity was evaluated by the measurement of halo diameters around bacterial colonies (in cm) relative to colony diameter on NBRIP (National Botanical Research Institute’s Phosphate) medium. n00132 had a significantly higher phosphate solubilization index (2.36 cm) than n00007, n00008, n00010, n00011, n00016, n00064, n00069, n00082, n00128, n00170 and n00172. **(B)** Indoleacetic acid (IAA) production by phosphate solubilizing bacteria (PSB). *E. coli* strain OP50 was used as positive control for IAA production. n00132 produced a significantly higher amount of IAA than other PSB isolates, except for n00170. **(C)** Gibberellic acid (GA) production by phosphate solubilizing bacteria (PSB). *E. coli* strain OP50 used as negative control for GA production. PSB isolates n00014-1, n00014-2, n00015, n00064, n00069, n00078 and n00080 were not able to produce GA as they were not different from the negative control. * In all three figures statistical significance was determined by *t*-test. *: *p* < 0.05. Error bars show standard errors.

We next characterized these 25 phosphate solubilizing isolates for other possible plant growth promoting traits, including nutrient sequestration and hormone production (3). Seventeen of the 25 (68%) PSB isolates were considered nitrogen fixing microorganisms that could also contribute to plant growth (Supplementary Fig. S2 (E) and Supplementary Table S2). Only one isolate (n00078) was able to solubilize zinc sulfate (Supplementary Fig. S2 (D)). Of the phytohormones tested, all 25 PSB isolates had the ability of indoleacetic acid (IAA) production, and among them, isolate n00132 produced the highest amount of IAA (10.8 μg mL^-1^; *p* < 0.05), two to three times more than other isolates (Fig. 1 (B)). Gibberellic acid production was observed in 72% (18/25) of the isolates (Fig. 1 (C)).

High soil salinity negatively affects the ability of plant roots to take up water and bacteria that can tolerate high concentrations of NaCl were previously shown to protect plants from water deficiency stress (56). Therefore, all 25 PSB isolates were tested for salt tolerance on LB medium with different concentrations of NaCl (2%, 4%, 6% and 8%). All 25 PSB were able to tolerate at least 2% NaCl and 80% (20/25) of them grew on LB with 4% NaCl (Supplementary Table S4). Only one bacterium (n00014-1) grew on LB with 6% NaCl and none of the PSB were able to tolerate 8% NaCl.

Ethylene is a phytohormone that plant roots produce for developmental processes such as xylem formation and to regulate stress responses (57). When plants are under stress, they often emit excessive ethylene, which has a negative effect on plant development. Bacteria that produce 1-aminocyclopropane-1-carboxylic acid (ACC) deaminase convert ACC, a rate limiting precursor for ethylene production, to ammonia and α-ketobutyrate, which can protect plants from excessive amounts of ethylene (58). In this study, PSB that produced ACC deaminase enzyme were examined based on their ability to grow on DF minimal salt medium supplemented with 3 mM ACC as sole nitrogen source (Supplementary Fig. S2 (F) and Supplementary Table S5). 84% (21/25) of the isolates had ACC deaminase activity that might protect plants from toxic ethylene buildup (Table S3).

PSB isolates were next tested for antifungal activity, lipase production, protease activity (casein and gelatin hydrolyzing), cellulase and ammonia production, all of which can contribute to bacterial defense mechanisms against plant pathogens (27). PSB isolates were also tested for siderophore production, which is a low molecular mass compound with high affinity to Fe^3+^ and facilitates iron uptake by plants (59). Results for these indirect PGPB assays are summarized in Supplementary Table S6. Based on the results, 68% (17/25) of the PSB isolates were able to facilitate iron uptake in plants and produce lipase enzyme, and 84% (21/25) of phosphate solubilizing bacteria were able to produce cellulase enzymes. 56% (14/25) of the phosphate solubilizing isolates were able to liquify gelatin, among which 71% (10/14) strongly liquified it and 29% (4/14) slightly liquified it. Only 8% (2/25) of the phosphate solubilizing isolates were able to hydrolyze casein. 84% (21/25) of the PSB isolates were able to inhibit fungal pathogen growth. All 25 PSB isolates were able to produce ammonia. These results showed that all 25 bacterial isolates tested had some ability to indirectly improve plant growth through pathogen control.

### Whole genome sequencing and phenotypic analysis of five phosphate solubilizing bacteria

Five bacterial isolates that were positive for phosphate solubilization, IAA and gibberellic acid production, and for nitrogen fixation were selected for whole genome sequencing. Genome assembly statistics and taxonomic assignments for the five isolates are shown in Table 1. Three of the isolates (n00163, n00167, n00172) were members of the Gram-positive *Paenibacillaceae* family, while the others belonged to the Gram-negative Proteobacteria phylum (Fig. 2). Three of the five isolates (n00163, n00167, n00170) fell near or below the 95% average nucleotide identity (ANI) threshold that is sometimes used for species delineation (40, 60, 61), indicating that these isolates are potentially new species. The other two isolates were new strains of *Pseudomonas mosselii* and *Paenibacillus graminis* (Table 1). All five of the isolates had genes predicted for synthesis of the phytohormone auxin (IAA), for ammonia assimilation, and for phosphate metabolism. These genome annotations provide the genotypes to accompany their plant growth promoting phenotypes.

**Table 1.**
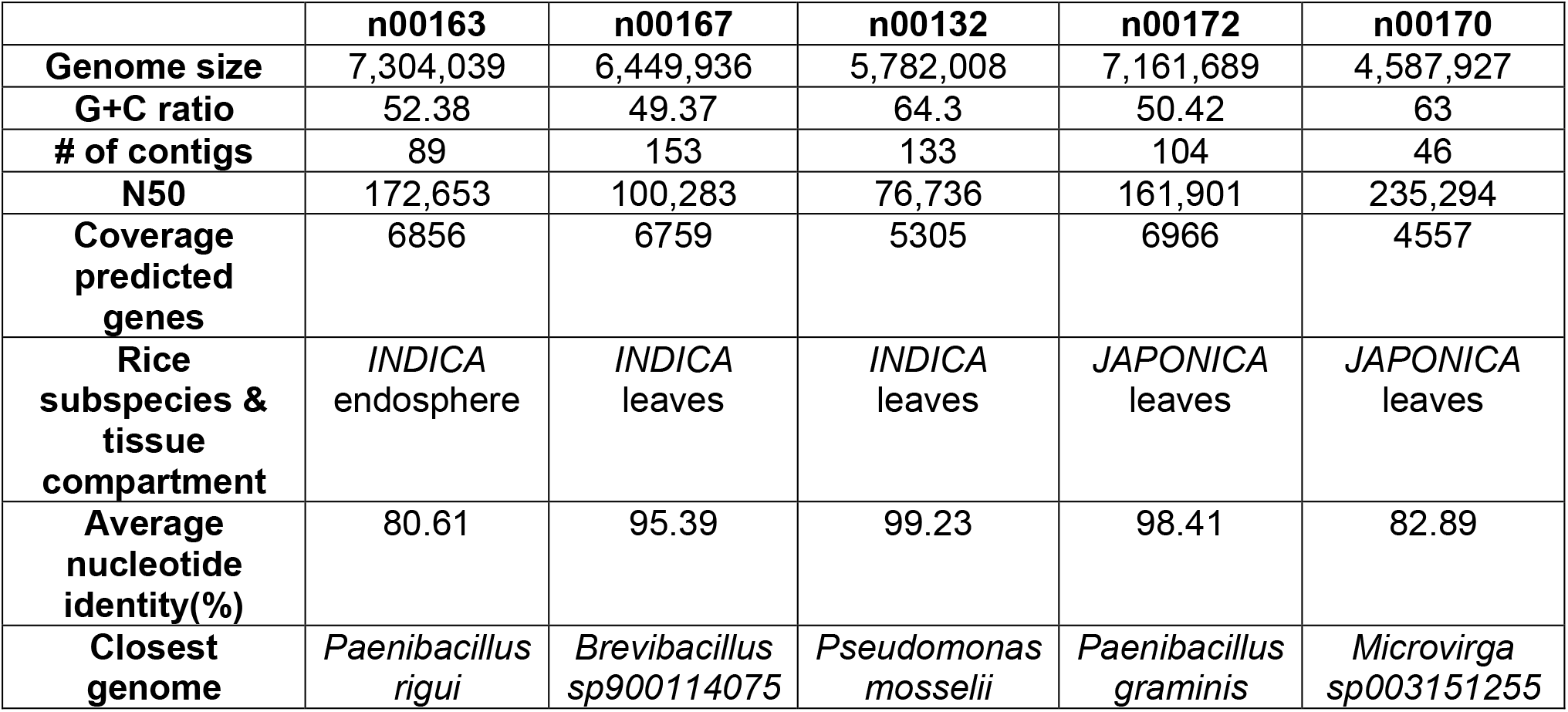
Feature types and assemble Reads with SPAdes - v3.13.0 results (QUAST analysis) of sequenced genomes and genome Taxonomy Database (GTDB) analysis results of phosphate solubilizing bacteria isolated from different rice subspecies and tissues

**FIG 2.**
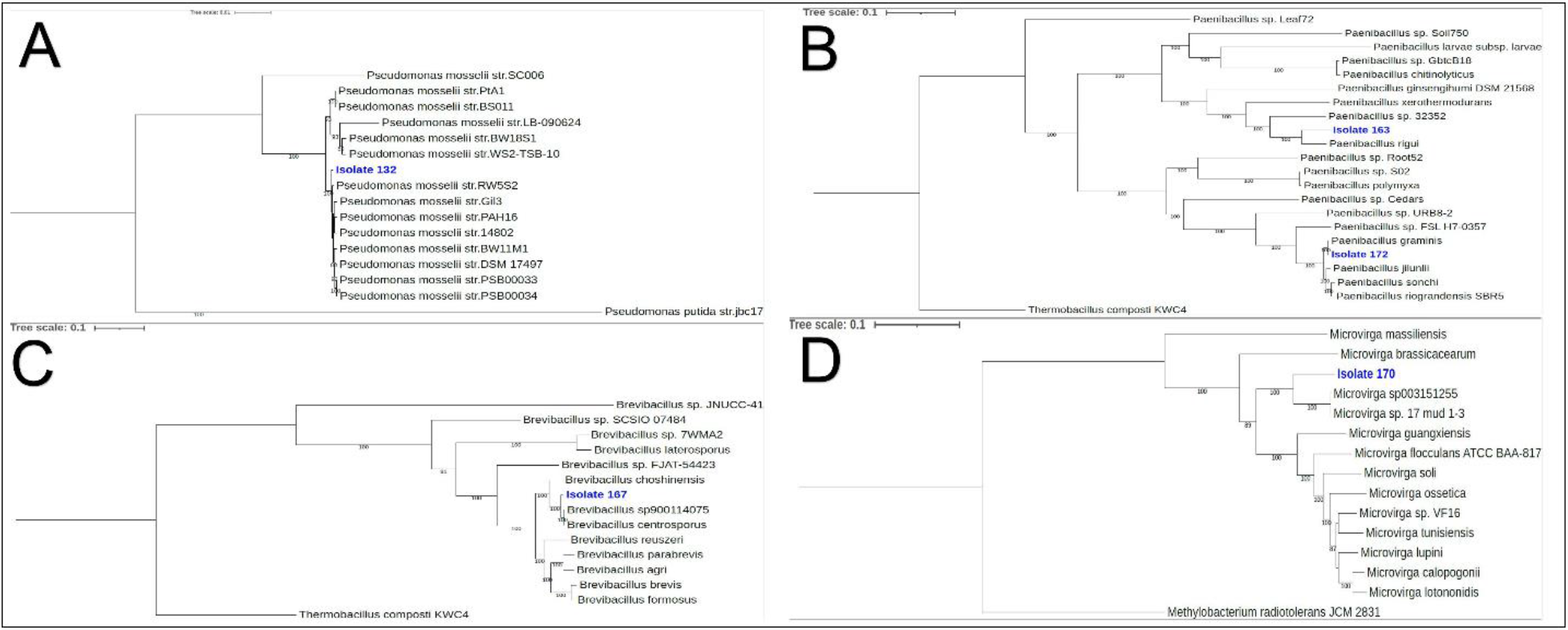
Phylogenetic trees based on whole genome sequences of five bacteria isolated from different rice tissue compartments. **(A)** *Pseudomonas mosselii*, isolated from *INDICA* leaves. **(B)** *Paenibacillus rigui*, isolated from *INDICA* endosphere and *Paenibacillus graminis*, isolated from *JAPONICA* leaves. **(C)** *Brevibacillus sp900114075*, isolated from *INDICA* leaves. **(D)** *Microvirga sp003151255*, isolated from *JAPONICA* leaves.

Gen III microplates (Biolog) were used to characterize the metabolic phenotypes of the five sequenced PSB. *Agrobacterium tumefaciens*, a Gram-negative plant pathogen producing crown gall tumors (62), was used as a comparison because of its well-characterized ability to use plant metabolites as nutrients (63). Of the five isolates tested, two (n00163 and n00167) shared similar phenotypic carbon source usage characteristics (Supplementary Table S7). Of the different metabolic phenotypes, methyl pyruvate is a plant chemoeffector (64) and glycerol is a leaf exudate as a result of carbon dioxide fixation in photosynthesis (65). Glycerol was not used by n00132, while methyl pyruvate was not used by n00132 and n00172. Interestingly, n00172 was the only bacterium able to utilize stachyose (raffinose family of oligosaccharides in plants) (66). All five sequenced PSB isolates shared the same phenotypic chemical resistance against pH 6, 1% NaCl, 1% sodium lactate, guanidine HCl, tetrazolium blue, nalidixic acid, lithium chloride, potassium tellurite and aztreonam. Only one bacterium (n00172) showed resistance against Rifamycin SV and sodium butyrate (Supplementary Table S8).

### Antibacterial activity of the sequenced PSB against each other

Overlay-agar and cross streak assays were done to determine whether the five sequenced PSB were compatible with each other. Plants derive greater benefits from the mixture (consortium) of PGPB than one bacterium (67). Therefore, it is important to investigate whether these five PSB would compete, act neutrally, or synergistically to improve rice growth. Based on these assays, only isolate n00172 competitively inhibited the growth of the other four PSB (Supplementary Fig. S2 (A & B)). This indicates that n00172 (*Paenibacillus graminis*) produced inhibitory chemicals that did not allow other bacteria to grow in its close proximity (Supplementary Table S9). Analysis of n00172 genome through the program antiSMASH (68) revealed this bacterium has several biosynthetic gene clusters that are predicted to have antibiotic activity (Supplementary Table S10) that may be responsible for competitive inhibition of other species.

### Influence of sequenced PSB on rice plant growth and development

To determine whether the five sequenced PSB act as PGPB, rice seeds from *INDICA* and *JAPONICA* accessions were inoculated individually with one of the five PSB, and the shoot and root development (length) and biomass (dry weight) of two-week old plants were compared to uninoculated control plants. Control plants were from uninoculated seeds that were soaked in KCl solution without any bacteria. *JAPONICA* plants inoculated with n00132 (*Pseudomonas mosselii*), n00170 (*Microvirga* sp.), and n00172 (*Paenibacillus graminis*) had significantly longer roots than uninoculated controls (Fig. 3) (*p* < 0.001). In addition to root length, *JAPONICA* plants inoculated with n00132 (*P. mosselii*), n00163 (*Paenibacillus rigui*), and n00172 (*P*. *graminis*) also had significantly longer shoots than controls (Fig. 3) (*p* < 0.05 to 0.001). Taken together, this shows that those four species are PGPB for the *JAPONICA* subspecies of rice.

**FIG 3.**
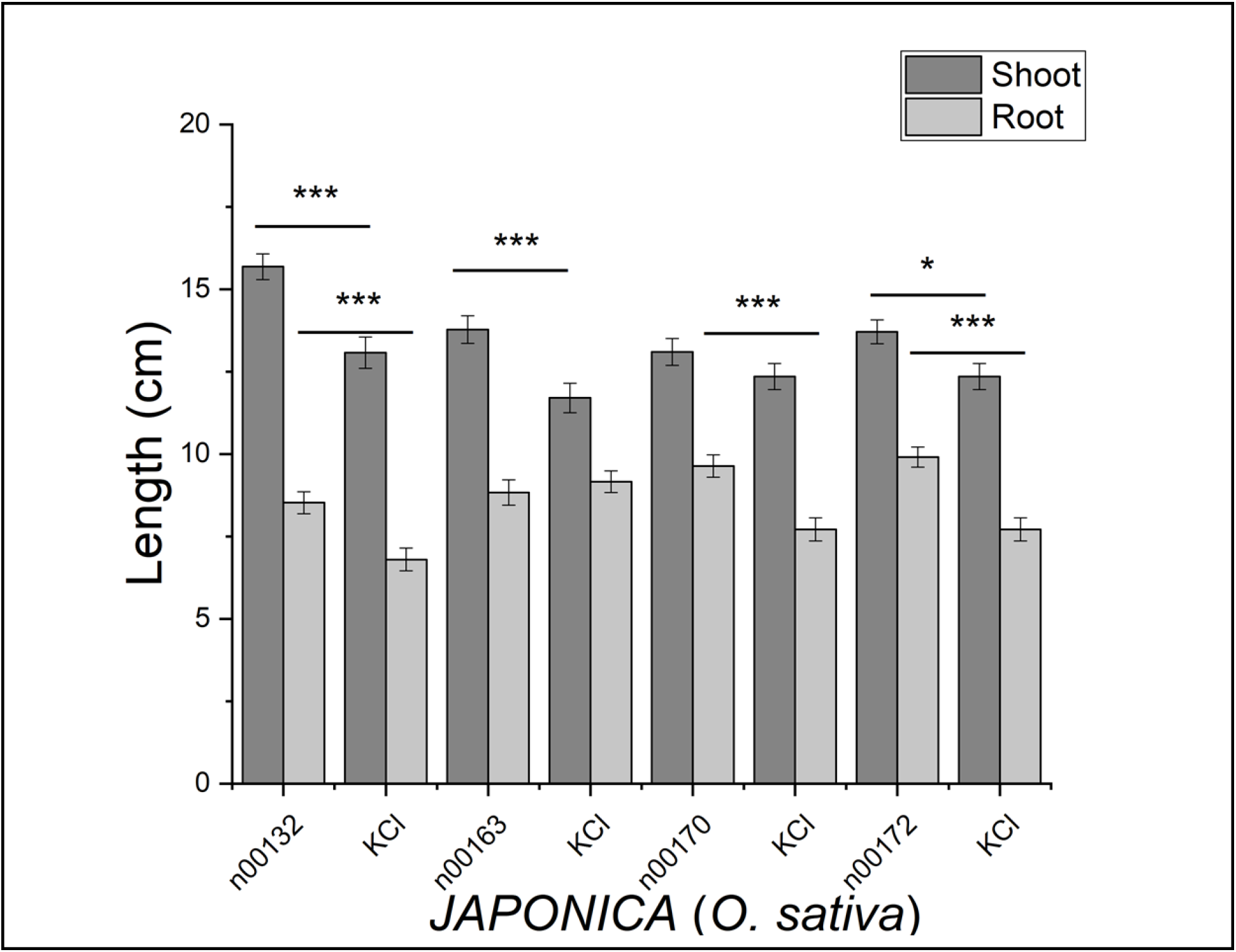
Root and shoot lengths of 2-week-old *JAPONICA* plants inoculated with bacteria compared to uninoculated control plants (KCl). n00132, plants inoculated with *Pseudomonas mosselii*; n00163, plants inoculated with *Paenibacillus rigui*; n00170, plants inoculated with *Microvirga* sp.; n00172, plants inoculated with *Paenibacillus graminis*. Statistical significance was determined by *t*-tests. *: *p* < 0.05; **: *p* < 0.01; ***: *p* < 0.001

In contrast, *INDICA* plants inoculated with the five bacteria did not have siginificantly improved root or shoot growth (Supplementary Fig. S3). Surprisingly, *INDICA* plants inoculated with n00170 (*Microvirga* sp.) and n00172 (*P. graminis*) had significantly shorter roots than uninoculated control plants (Fig. 4) (*p* < 0.01 to 0.001). This shows that those two isolates were not PGPB for the *INDICA* subspecies of rice.

**FIG 4.**
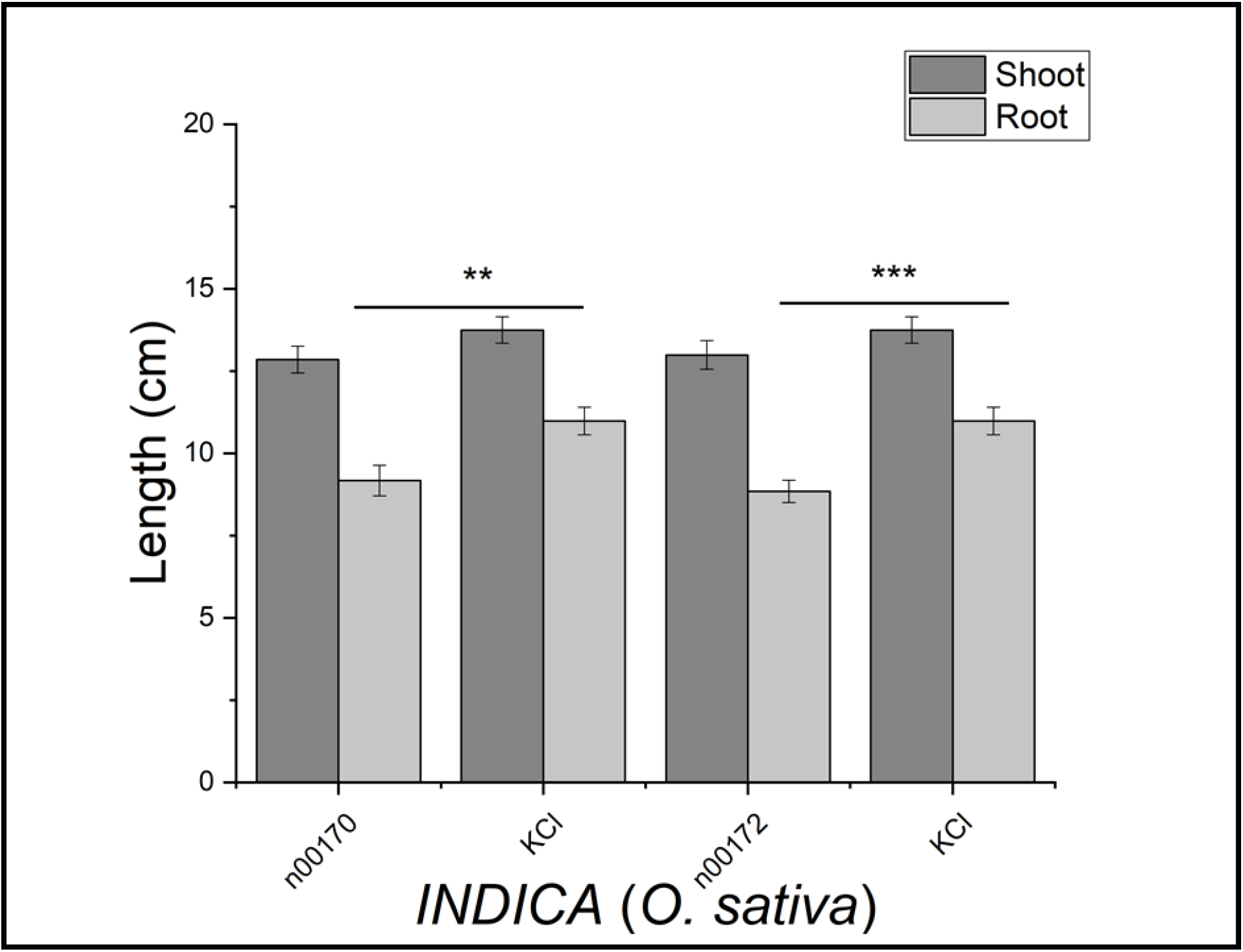
Root and shoot lengths of 2-week-old *INDICA* plants inoculated with bacteria compared to uninoculated control plants (KCl). n00170, plants inoculated with *Microvirga* sp.; n00172, plants inoculated with *Paenibacillus graminis*. Statistical significance was determined by *t*-tests. *: *p* < 0.05; **: *p* < 0.01; ***: *p* < 0.001

Root/shoot ratio refers to the biomass that is growing underground and reflects the ability of plants to take up nutrients. The rationale for assessing this is that the higher this ratio, the greater the ability of plants to take up nutrients, which correlates with increased stress tolerance (51). Our measurements showed that *JAPONICA* plants inoculated with n00132 (*P. mosselii*) had significantly higher dry weight root/shoot ratios (*p* = 4.2×10^-2^), suggesting that this species helps with nutrient uptake (Fig. 5 (A)). Moreover, *JAPONICA* plants inoculated with n00132 (*P. mosselii*) and n00172 (*P*. *graminis*) had higher root dry weights than control plants and plants inoculated with other bacteria (Fig. 6 (A) (*p* < 0.05).

**FIG 5.**
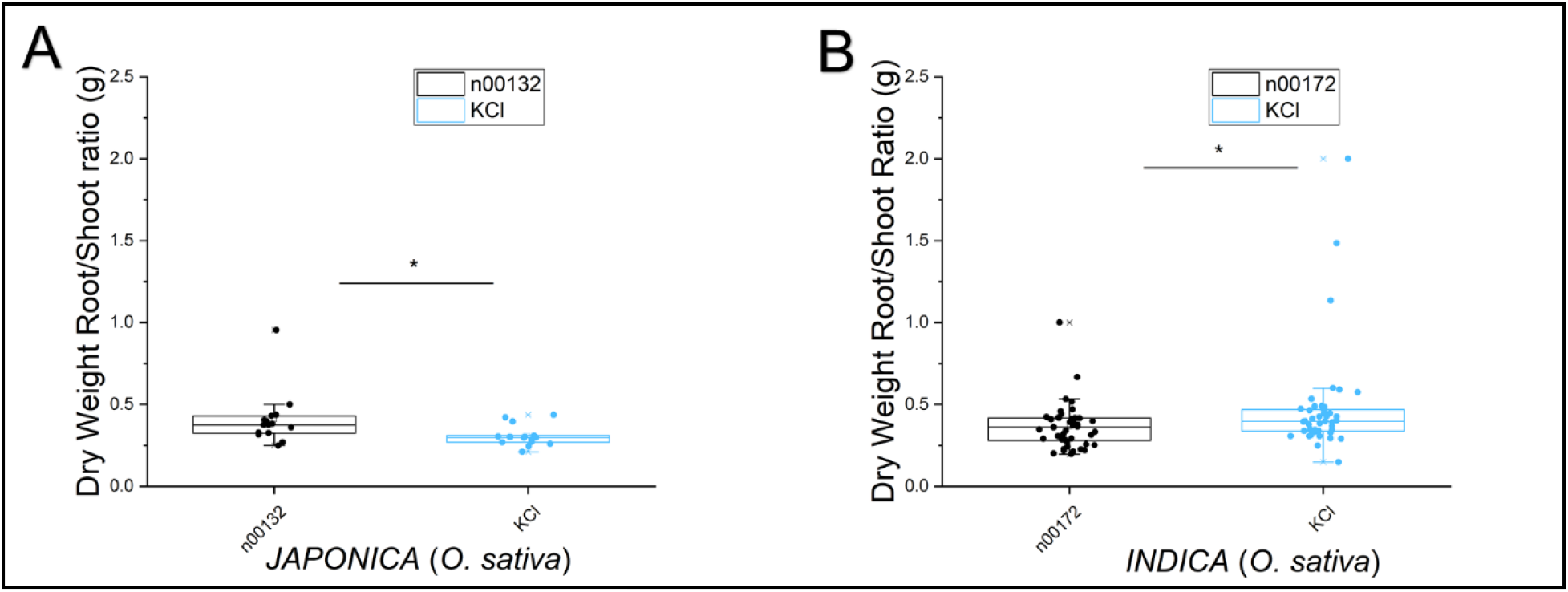
Dry weight root/shoot ratios of 2-week-old *JAPONICA* and *INDICA* plants compared to uninoculated control plants (KCl). **(A)** n00132; *JAPONICA* plants inoculated with *Pseudomonas mosselii*. **(B)** n00172, *INDICA* plants inoculated with *Paenibacillus graminis*. Each dot represents an individual plant. Statistical significance was determined by *t*-test. *: *p* < 0.05

**FIG 6.**
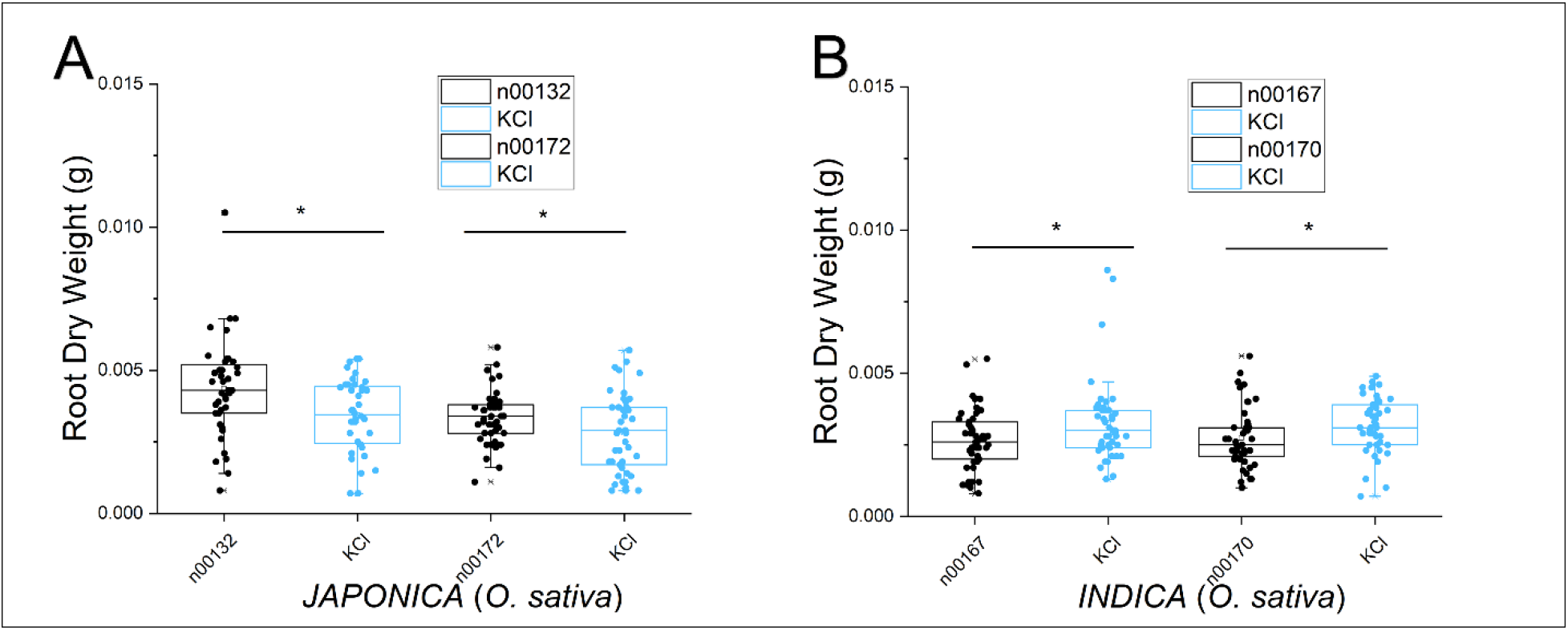
Dry weight of 2-week-old *JAPONICA* and *INDICA* roots inoculated with bacteria compared to uninoculated control plants (KCl). **(A)** n00132 and n00172, *JAPONICA* plants inoculated with *Pseudomonas mosselii* and *Paenibacillus graminis*, respectively. **(B)** n00170 and n00167, *INDICA* plants inoculated with *Brevibacillus* sp. and *Microvirga* sp., respectively. Each dot represents an individual plant. Statistical significance was determined by *t*-test. *: *p* < 0.05

In contrast, *INDICA* plants inoculated with the five bacteria did not have significantly improved root or shoot dry weights or root/shoot ratios than uninoculated control plants (Supplementary Fig. S4). Contrary to the results obtained with *JAPONICA* plants, *INDICA* plants inoculated with n00172 (*P. graminis*) had significantly lower root/shoot ratios than control plants (Fig. 5 (B)) (*p* < 0.05). Root dry weights of *INDICA* plants inoculated with *Brevibacillus* sp. and *Microvirga* sp. were also significantly lower than those of control plants (Fig. 6 (B) (*p* < 0.05).

## Discussion

To identify plant growth promoting bacteria (PGPB) from rice, we isolated 140 bacteria residing in or on different tissues of plants representing both the *INDICA* and *JAPONICA* subspecies of rice and screened for phosphate solubilizing activity. 25 of the isolates were phosphate solubilizing bacteria (PSB) that were also able to produce indole acetic acid, an important plant growth hormone. Based on other direct and indirect PGPB assays, five PSB were selected for whole genome sequencing. The five PSB species were closely related to *Paenibacillus rigui* (n00163, isolated from *INDICA* endosphere), *Brevibacillus sp.* (n00167, isolated from *INDICA* leaves), *Pseudomonas mosselii* (n00132, isolated from *INDICA* leaves), *Paenibacillus graminis* (n00172, isolated from *JAPONICA* leaves), and *Microvirga sp.* (n00170, isolated from *JAPONICA* leaves).

Although only four of the PSB improved *JAPONICA* subspecies growth and development, all of these bacterial genera were previously shown to function as PGPB (19, 62, 69, 70, 71). Some *Brevibacilllus* species are able to produce antifungal siderophores (72) while the *Microvirga* genus is a nodule legume endophyte that can improve plant growth by providing ammonia to plants (71). Some species of *Paenibacillus* have been isolated from plant rhizospheres, and *P. graminis* is a known nitrogen fixing bacterium (69, 73). *P. rigui* is one of the species that has been specifically isolated as an endophyte from *O. sativa* (74). *Pseudomonas* species are widely present in soils, and it is known that some strains can survive in different environmental conditions and tissues of eukaryotic hosts mostly through producing chemicals that protect them against pathogenic bacteria or fungi (62, 70). *P*. *mosselii* was previously shown to be a phosphate solubilizing bacterium (80), and our results are in agreement with this finding (Fig. 1 (A)). *P. mosselii* is also one of the most common PGPB that inhibits growth of plant pathogens such as *Agrobacterium tumefaciens* (62) and results from our study (cross-streak experiment) confirms this observation (Supplementary Fig. S5). In addition, *P. mosselii* is also known for improving plant growth and increasing sugar content of *Agave americana* L. (70), and our results are in agreement with some of these findings (Figs. 3 & 5). Therefore, this bacterium is a good candidate for promoting plant growth, and potentially also for protecting plants against stress (75).

Auxins such as IAA are an important class of plant growth hormones (phytohormones) (26), and auxin producing PSB might be PGPB by stimulating root growth and differentiation. The 25 PSB isolates were tested for IAA production, which was previously shown to act as a plant growth promoting phytohormone when produced by plant-associated bacteria (72, 76). Our results specifically showed that n00132 (*P. mosselii*) was the bacterium with the highest phosphate solubilizing index (2.36 cm; Fig. 1 (A)) and highest IAA production (10.81 μg mL^-1^; Fig. 1 (B)). In addition, this bacterium was the only one with positive results for all indirect PGPB pathways including siderophore, cellulase, lipase, protease, and ammonia production. Results from seed inoculations moreover showed that n00132 (*P. mosselii)* was the only bacterium that significantly increased root and shoot growth and the dry weight root/shoot ratio of *JAPONICA* rice plants. Other PSB that we isolated including n00163 (*P. rigui*), n00170 (*Microvirga* sp.), and n00172 (*P. graminis*) improved one or the other parameter. However, based on the genomic analysis, n00172 (*P. graminis*) and n00170 (*Microvirga* sp.) do not contain sequences for siderophore production, and this might be one of the reasons why these bacteria had a negative effect on root growth of *INDICA* plants, because bacterial siderophore has been shown to have a direct beneficial effect on plant growth (77) Also, *Brevibacillus* sp. does not contain denitrifying reductase gene clusters, which can contribute to root formation and development (78). Taken together, this might explain why among the five sequenced PSB, n00167 (*Brevibacillus* sp.) did not improve root growth of *JAPONICA* plants and negatively influenced root biomass (root dry weight) of *INDICA* plants.

Interestingly, the results of this study showed that n00132 (*P. mosselii*), n00170 (*Microvirga* sp.), n00172 (*P. graminis*), and n00163 (*P. rigui*) were able to increase root and/or shoot growth and root biomass of two-week-old *JAPONICA* plants but had no or a negative effect on the growth of two-week-old *INDICA* plants. To the best of our knowledge, this is the first work describing a positive effect of PSB that were isolated from both *INDICA* and *JAPONICA* accessions on only one subspecies of *O. sativa*. There are at least two possibilities for this observation. First, genetic differences between *JAPONICA* and *INDICA* plants might lead to differential response to bacterial metabolisms (79). It is reasonable to assume that PSB interact with and possibly regulate plant genes for growth and development. These genes could be differentially expressed between *JAPONICA* and *INDICA* or absent in one of the subspecies. Second, there might be differences in how genes/operons and/or metabolic pathways within the five PSB respond to different exudates and plant hormones. Further studies need be done to understand the mechanisms of how these bacteria respond differently to different rice varieties, and *vice versa*, how different rice varieties respond differently to these bacteria. While many studies have discovered the benefits of plant growth promoting bacteria (3, 15, 25, 74, 79), we demonstrate here that the benefits may not be universally applied. Even plant taxa as closely related as subspecies will respond differently to plant growth promoting bacteria, indicating the need to develop strain specificities between plants and bacteria.

## Acknowledgements

We would like to thank Dr. Stacia Peiffer for technical support and graduate student Huy Phan for help with some experiments. This research was funded in part by National Institute of Food and Agriculture-Agriculture and Food Research Initiative (NIFA-AFRI) grant number 2016-67013-24587 from the United States Department of Agriculture (USDA). The sequencing part of this work was supported by Marquette University Department of Biological Sciences educational funding.

---

**FIG S1.**
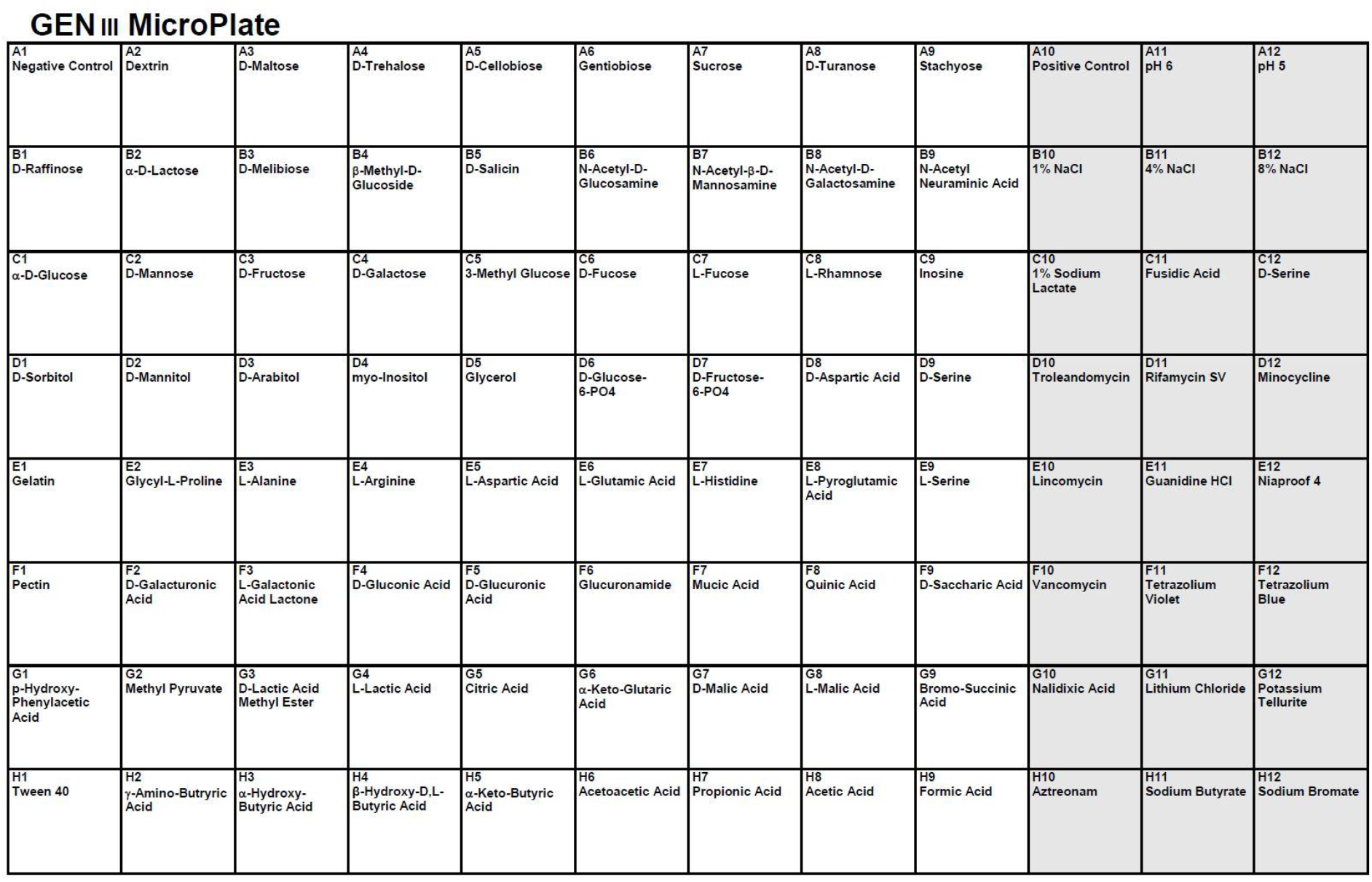
Layout of 71 carbon sources and 23 chemical sensitivity assays on the 96 well microplate.

**FIG S2.**
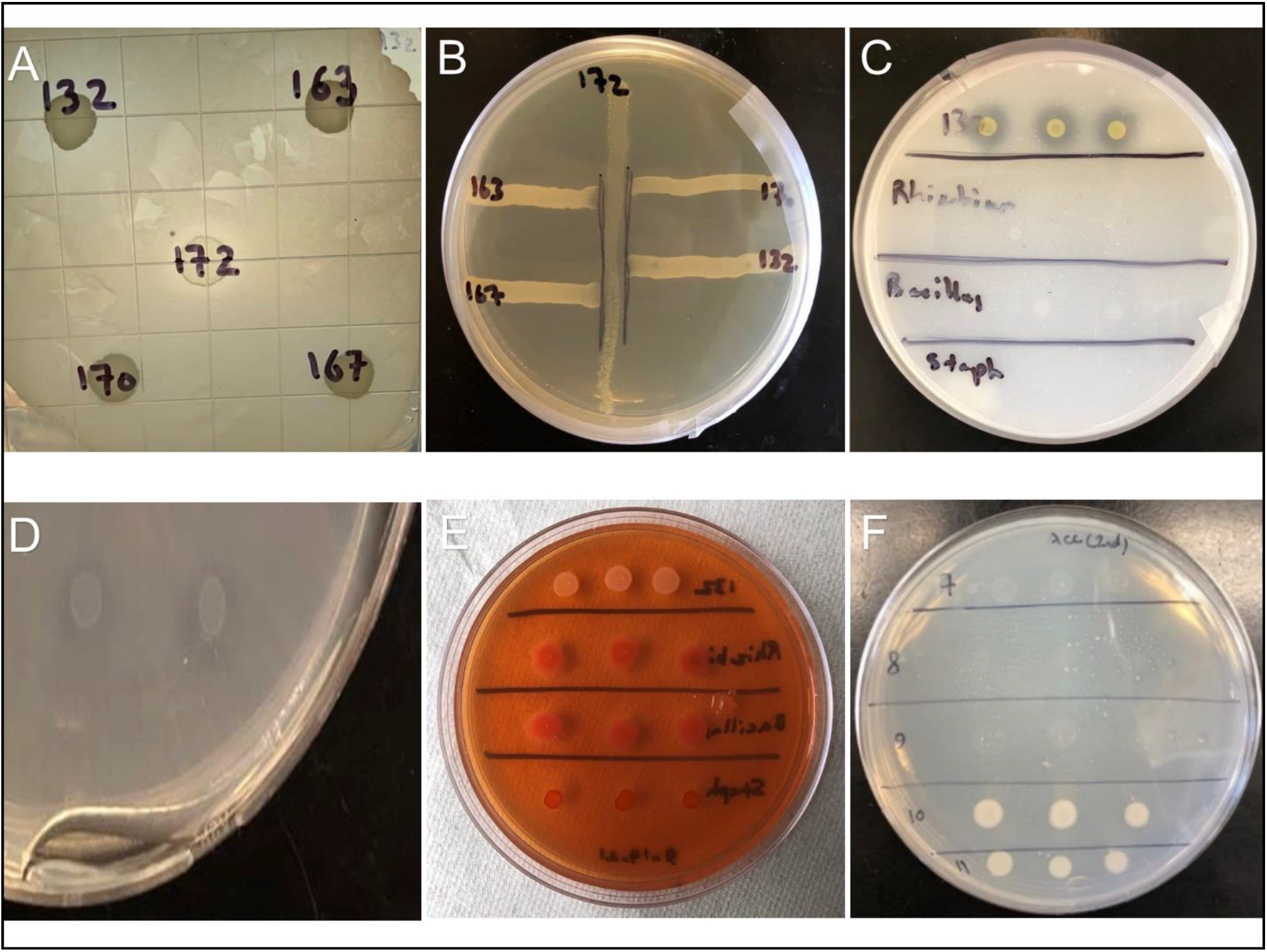
**(A)** Overlay agar assay. Halo around 172 (*Paenibacillus graminis*), showing growth inhibition when 132 (*Pseudomonas mosselii*) is overlaid. Halo around 172 was also observed, when the other three phosphate solubilizing bacteria (PSB) were overlaid: 163, *Paenibacillus rigui*; 167, *Brevibacillus sp900114075*; 170, *Microvirga sp003151255.* **(B)** Cross-streak assay. Growth inhibition of the other four PSB is shown when 172 is in the middle of the NB plate. **(C)** Bacterium 132 isolated from the *INDICA* phyllosphere has high phosphate solubilization ability as shown by strong halo formations (2.36 cm) around the bacterial colonies. *Rhizobacterium* sp., *Bacillus* sp., and *Staphylococcus aureus* are negative controls. **(D)** Bacterium n00078 isolated from *JAPONICA* root endosphere has zinc solubilization ability as shown by halo formation around the bacterial colony. **(E)** Bacteria with nitrogen fixation ability shown on CR-YMA plates. The lower the absorption of Congo Red dye, the higher the ability for nitrogen fixation. In this figure, one strong nitrogen fixing bacterium (top 3 colonies; *P. mosselii*) is along with *Rhizobacterium* sp. and *Bacillus* sp., as positive controls and *Staphylococcus aureus* as a negative control. **(F)** Phosphate solubilizing bacteria n00007 (7), n00008 (8), n00009 (9), n00010 (10), and n00011 (11) are shown with the ability of ACC deaminase production based on their growth on DF minimal salt medium supplemented with 3 mM ACC as sole nitrogen source.

**FIG S3.**
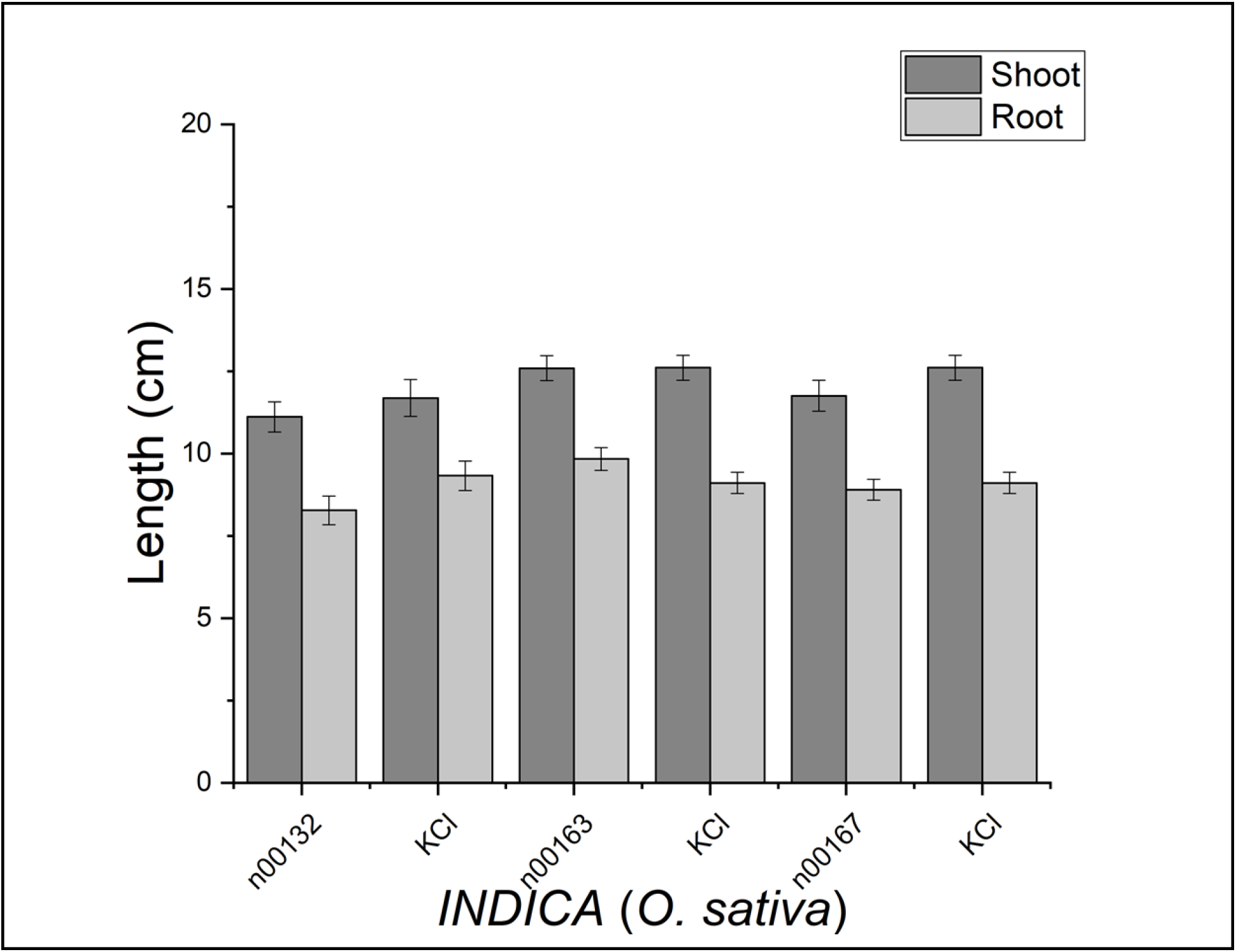
Root and shoot lengths of 2-week-old *INDICA* plants inoculated with bacteria compared to uninoculated control plants (KCl). n00132, plants inoculated with *Pseudomonas mosselii*; n00163, plants inoculated with *Paenibacillus rigui*; n00167, plants inoculated with *Brevibacillus* sp. No significant differences between inoculated and uninoculated plants were found.

**FIG S4.**
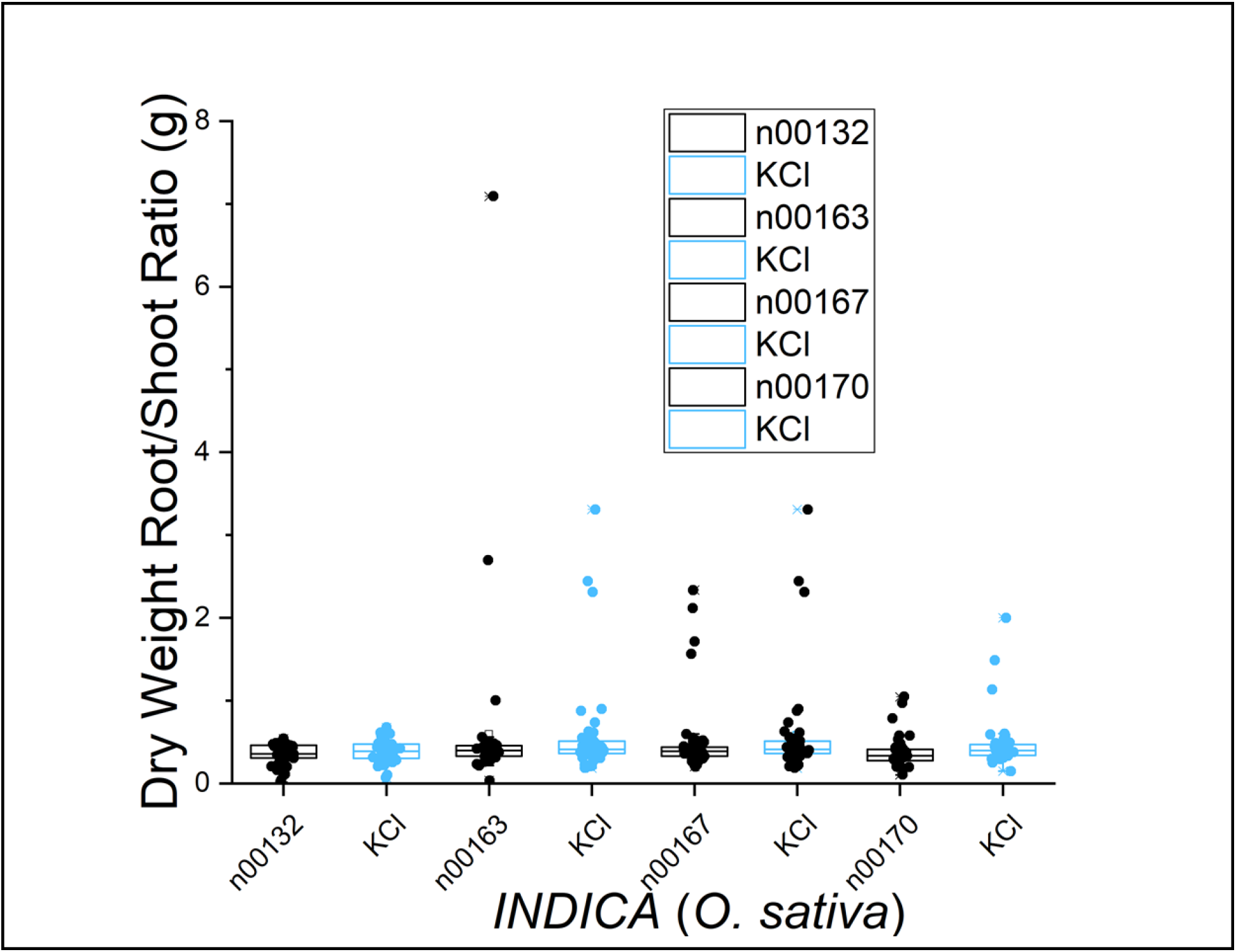
Dry weight root/shoot ratios of 2-week-old *INDICA* plants compared to uninoculated control plants (KCl). n00132, plants inoculated with *Pseudomonas mosselii*; n00163, plants inoculated with *Paenibacillus rigui;* n00167, plants inoculated with *Brevibacillus* sp.; n00170, plants inoculated with *Microvirga* sp. No significant differences between inoculated and uninoculated plants were found.

**FIG S5.**
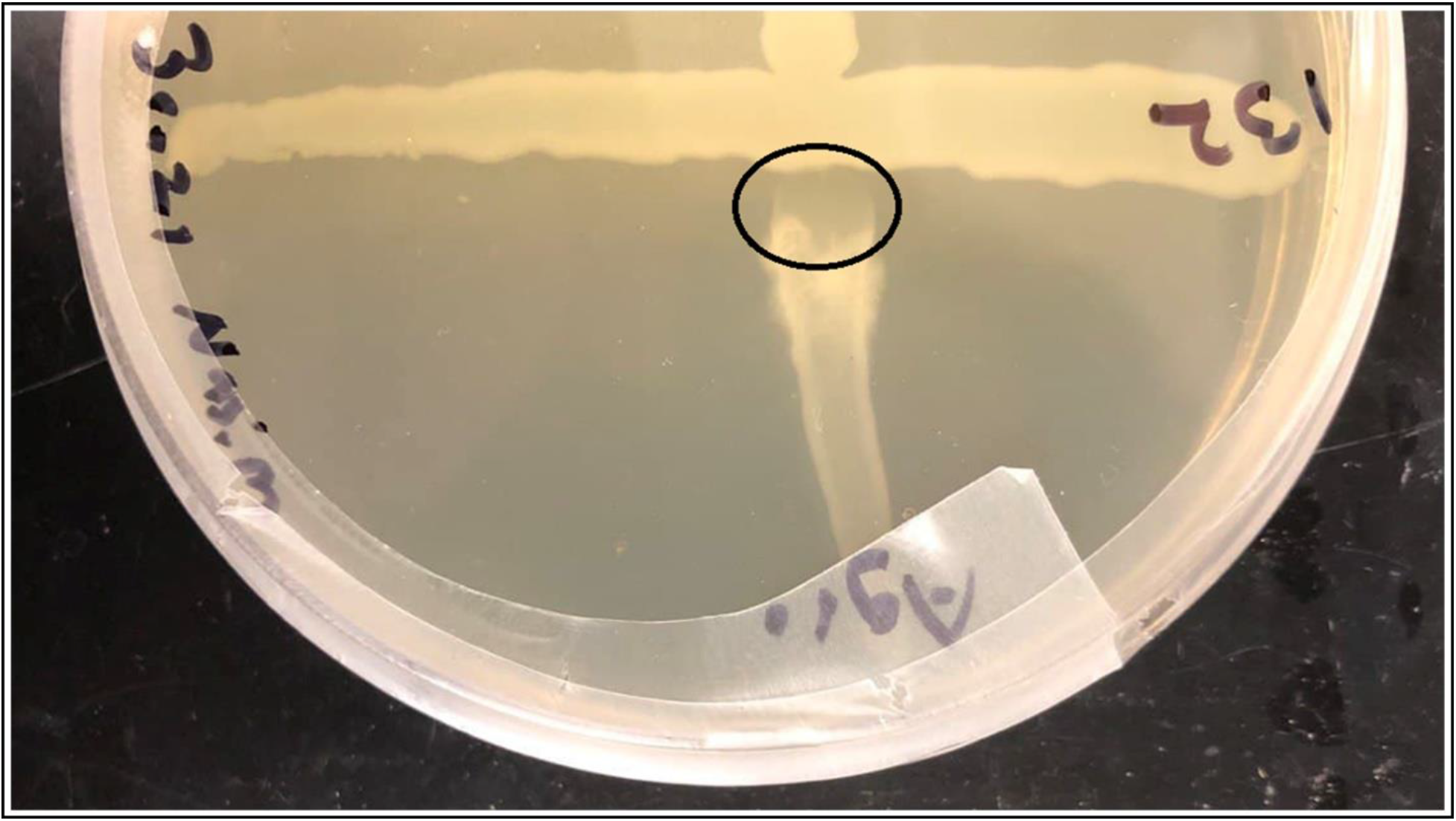
Cross-streak assay. Growth inhibition of the *Agrobacterium tumefaciens* is shown (circle) when 132 (*Pseudomonas mosselii*) is in the middle of the NB plate.

**Table S1.** List of culture-based plant growth promoting assays

| Direct mechanisms of PGPB assays | Reference(s) |
| --- | --- |
| Zinc solubilization | (21) |
| Nitrogen fixation | (22, 23) |
| Indoleacetic acid (IAA) production (phytohormone) | (24) |
| Gibberellic acid production (phytohormone) | (25, 26) |
| 1-aminocyclopropane-1-carboxylic acid (ACC) deaminase activity | (21) |
| Indirect mechanisms of PGPB assays | Reference(s) |
| Antifungal activity | (27, 28) |
| Lipase production | (56) |
| Casein and gelatin hydrolyzing (proteases) | (30) |
| Cellulase production | (81) |
| Siderophore production | (31) |
| Ammonia (NH<sub>3</sub>) production | (33) |

**Table S2.**
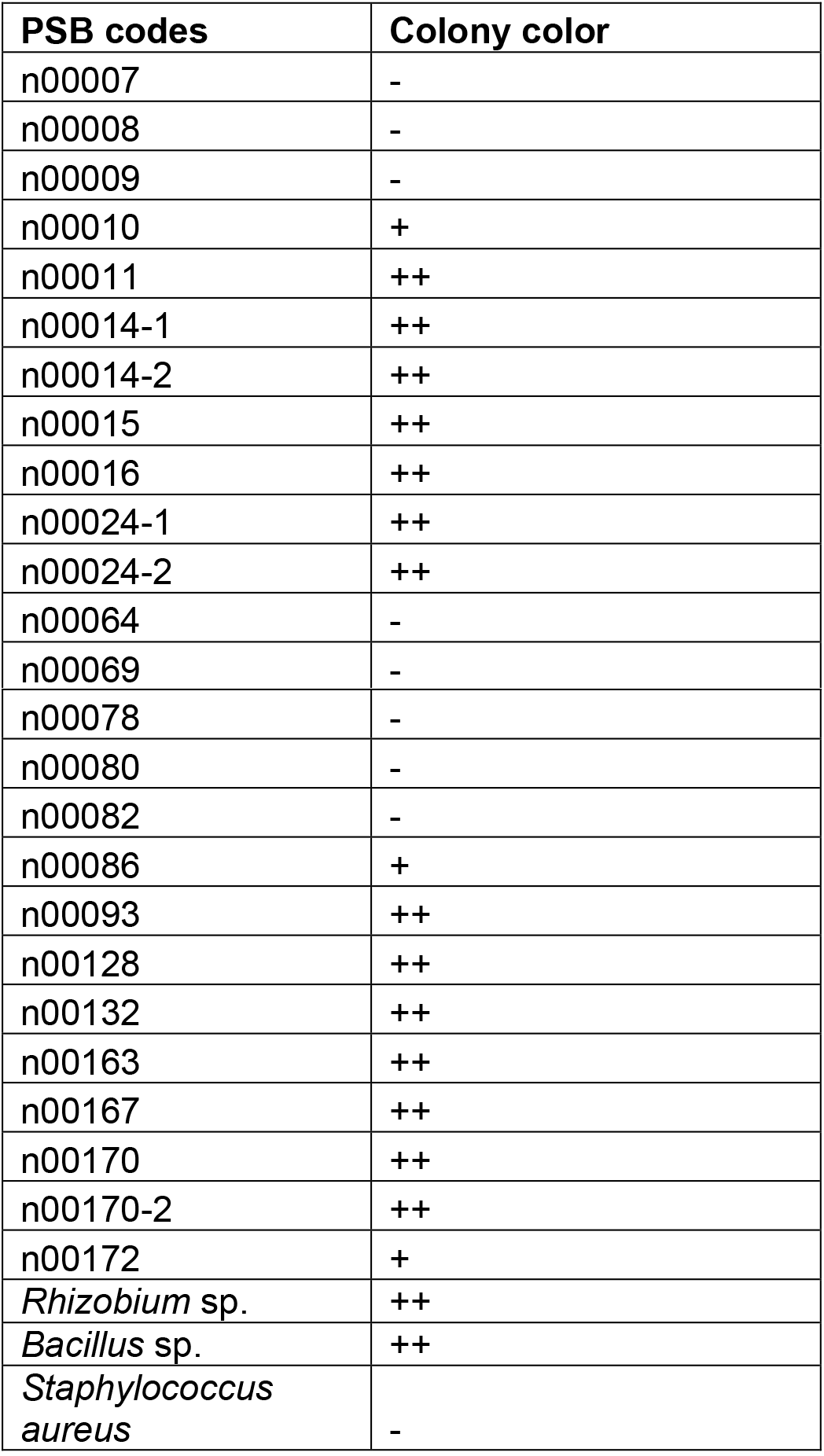
Phosphate solubilizing bacteria with nitrogen fixation ability on CR-YMA plates. ++ (Pale white-pink), + (White-pink), - (Strong pink-red). Bacterial colonies with colors ranging from white to pale white-pink were able to fix nitrogen due to cleavage of the azo bond (-N=N-) in Congo Red.

| PSB codes | Colony color |
| --- | --- |
| n00007 | - |
| n00008 | - |
| n00009 | - |
| n00010 | + |
| n00011 | ++ |
| n00014-1 | ++ |
| n00014-2 | ++ |
| n00015 | ++ |
| n00016 | ++ |
| n00024-1 | ++ |
| n00024-2 | ++ |
| n00064 | - |
| n00069 | - |
| n00078 | - |
| n00080 | - |
| n00082 | - |
| n00086 | + |
| n00093 | ++ |
| n00128 | ++ |
| n00132 | ++ |
| n00163 | ++ |
| n00167 | ++ |
| n00170 | ++ |
| n00170-2 | ++ |
| n00172 | + |
| *Rhizobium* sp. | ++ |
| *Bacillus* sp. | ++ |
| *Staphylococcus aureus* | - |

**Table S3.**
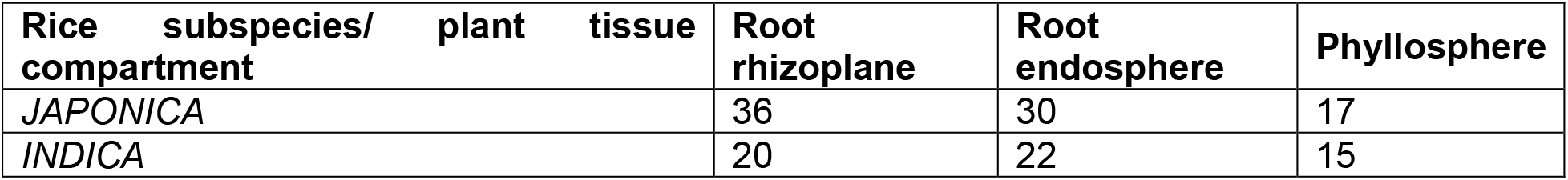
Number of bacterial isolates from two accessions, Krasnodarskij 3352 and Carolino 164, representing the *JAPONICA* and *INDICA* subspecies of rice (*Oryza sativa* L.), respectively.

**Table S4.** Phenotypic characteristics of isolated bacteria from *INDICA* and *JAPONICA* rhizoplane, endosphere, and phyllosphere.

| *INDICA*<br>rhizoplane<br>bacteria | Form | Elevation | Color | Size | Margin | Cell morphology | G+/G- |
| --- | --- | --- | --- | --- | --- | --- | --- |
| n00001 | Circular | Slightly raised | Yellow | M | Entire | Cocci | G+ |
| n00002 | Circular | Slightly raised | Milk white | M | Entire | Cocci | G+ |
| n00003 | Irregular | Flat | Milk white | L | Undulate | Rod-shape | G- |
| n00004 | Circular | Slightly raised | Milk white | S | Entire | Cocci | G+ |
| n00005 | Circular | Slightly raised | Milk white | S | Entire | Cocci | G+ |
| n00006 | Circular | Slightly raised | Milk white | M | Entire | Cocci | G+ |
| n00028 | Circular | Raised | Milk white | M | Entire | Cocci | G+ |
| n00029 | Circular | Raised | Yellow | S | Entire | Cocci | G+ |
| n00030 | Circular | Raised | Milk white | M | Entire | Cocci | G+ |
| n00031 | Circular | Raised | Yellow | S | Entire | Cocci | G+ |
| n00032 | Circular | Flat | Milk white | S | Entire | Cocci | G+ |
| n00033 | Circular | Raised | Milk white | M | Entire | Rod-shape | G- |
| n00092 | Circular | Raised | Yellow | L | Entire | Cocci | G+ |
| n00093 | Circular | Raised | White | S | Entire | Cocci | G+ |
| n00094 | Circular | Raised | Milk White | M | Entire | Rod-shape | G+ |
| n00111 | Circular | Raised | Yellow | S | Entire | Cocci | G+ |
| n00112 | Circular | Umbonate | White | L | Undulate | Rod-shape | G+ |
| n00113 | Circular | Raised | Yellow | S | Entire | Cocci | G+ |
| n00115 | Circular | Raised | Milk White | S | Entire | Cocci | G+ |
| n00116 | Circular | Raised | Yellow | S | Entire | Cocci | G+ |
| **INDICA**<br>**endosphere**<br>**bacteria** | **Form** | **Elevation** | **Color** | **Size** | **Margin** | **Cell**<br>**morphology** | **G+/G-** |
| n00012 | Circular | Flat | Milk white | S | Slightly undulate | Cocci | G- |
| n00013 | Circular | Slightly raised | Milk white | M | Undulate | Rod-shape | G+ |
| n00014 | Circular | Raised | Milk white | S | Filiform | Rod-shape | G+ |
| n00015 | Circular | Slightly raised | Milk white | M | Undulate | Cocci | G+ |
| n00041 | Circular | Slightly Raised | Milk white | M | Lobate | Rod-shape | G+ |
| n00042 | Circular | Slightly Raised | Milk white | L | Undulate | Rod-shape | G+ |
| n00043 | Circular | Flat | Milk white | M | Undulate | Cocci | G+ |
| n00044 | Circular | Raised | White | M | Lobate | Cocci | G+ |
| n00044-1 | Circular | Raised | White | S | Entire | Rod-shape | G+ |
| n00044-2 | Circular | Raised | Milk white | M | Undulate | Diplococci | G+ |
| n00044-3 | Circular | Raised | Milk white | M | Undulate | Rod-shape | G+ |
| n00045 | Circular | Flat | Milk white | M | Undulate | Cocci | G+ |
| n00070 | Irregular | Flat | Cream | L | Filiform | Rod-shape | G+ |
| n00106 | Circular | Raised | Yellow | M | Entire | Cocci | G+ |
| n00107 | Circular | Raised | Milk White | M | Entire | Cocci | G+ |
| n00108 | Circular | Raised | Yellow | S | Entire | Cocci | G+ |
| n00129 | Circular | Flat | Transparent | S | Entire | Cocci | G+ |
| n00130 | Circular | Slightly Raised | Milk White | S | Entire | Rod-shape | G+ |
| n00131 | Circular | Slightly Raised | Milk White | S | Entire | Cocci | G+ |
| n00147 | Circular | Flat | Milk White | M | Lobate | Rod-shape | G+ |
| n00163 | Circular | Raised | White | S | Entire | Rod-shape | G+ |
| n00173 | Irregular | Slightly Raised | Milk White | S | Lobate | Rod-shape | G+ |
| **INDICA**<br>**phyllosphere**<br>**bacteria** | **Form** | **Elevation** | **Color** | **Size** | **Margin** | **Cell**<br>**morphology** | **G+/G-** |
| n00022 | Irregular | Raised | Milk white | M | Undulate | Rod-shape | G+ |
| n00022-1 | Irregular | Raised | Milk white | S | Undulate | Cocci | G+ |
| n00023 | Irregular | Raised | Milk white | M | Lobate | Rod-shape | G+ |
| n00024 | insufficient growth | - | Milk white | M | - | Cocci | G+ |
| n00025 | Irregular | Raised | Milk white | M | Undulate | Cocci | G+ |
| n00025-1 | Circular | Slightly raised | Milk white | S | Entire | Cocci | G- |
| n00048 | Circular | Raised | Milk white | M | Lobate | Cocci | G+ |
| n00049 | Irregular | Slightly Raised | Milk white | L | Undulate | Chain Cocci | G+ |
| n00087 | Irregular | Flat | Cream | L | Filiform | Rod-shape | G+ |
| n00109 | Circular | Raised | Yellow | S | Entire | Diplo cocci | G+ |
| n00132 | Circular | Slightly Raised | Milk White | S | Undulate | Rod-shape | G- |
| n00157 | Circular | Raised | White | M | Entire | Diplo cocci | G+ |
| n00158 | Circular | Umbonate | Milk White | M | Undulate | Rod-shape | G+ |
| n00166 | Circular | Flat | Milk White | S | Entire | Cocci | G+ |
| n00167 | Circular | Flat | White | S | Entire | Rod-shape | G+ |
| **JAPONICA rhizoplane bacteria** | **Form** | **Elevation** | **Color** | **Size** | **Margin** | **Cell morphology** | **G+/G-** |
| n00007 | Irregular | Umbonate | Milk white | S | Filiform | Rod-shape | G- |
| n00008 | Irregular | Slightly raised | Milk white | M | Undulate | Cocci | G+ |
| n00009 | Irregular | Flat | Milk white | M | Undulate | Rod-shape | G- |
| n00010 | Circular | Slightly raised | Milk white | L | Entire | Cocci | G+ |
| n00011 | Circular | Slightly raised | Milk white | M | Filiform | Rod-shape | G- |
| n00034 | Circular | Raised | Yellow | L | Entire | Cocci | G- |
| n00035 | Circular | Raised | Milk white | S | Entire | Cocci | G+ |
| n00036 | Circular | Raised | Yellow | M | Entire | Cocci | G+ |
| n00039 | Circular | Raised | Yellow | M | Entire | Rod-shape | G+ |
| n00056 | Circular | Raised | Milk White | S | Entire | Rod-shape | G+ |
| n00057 | Circular | Raised | Milk White | M | Entire | Rod-shape | G+ |
| n00058 | Circular | Slightly Raised | Yellow | S | Entire | Rod-shape | G+ |
| n00059 | Circular | Slightly Raised | White | S | Entire | Rod-shape | G+ |
| n00060 | Circular | Slightly Raised | Milk White | S | Entire | Cocci | G+ |
| n00061 | Circular | Slightly Raised | Milk White | S | Entire | Rod-shape | G+ |
| n00062 | Circular | Crateriform | Red | M | Entire | Rod-shape | G+ |
| n00063 | Circular | Raised | Milk White | M | Entire | Rod-shape | G- |
| n00064 | Circular | Flat | Milk White | L | Undulate | Rod-shape | G+ |
| n00097 | Circular | Slightly Raised | Milk White | S | Entire | Rod-shape | G+ |
| n00098 | Circular | Flat | Yellow | S | Entire | Chain cocci | G+ |
| n00099 | Circular | Slightly Raised | Milk White | S | Entire | Cocci | G+ |
| n00101 | Circular | Raised | White | M | Entire | Cocci | G+ |
| n00102 | Circular | Flat | Transparent | S | Entire | Cocci | G+ |
| n00103 | Circular | Raised | Milk White | M | Entire | Rod-shape | G+ |
| n00104 | Circular | Raised | Yellow | L | Entire | Cocci | G+ |
| n00105 | Circular | Raised | White | S | Entire | Cocci | G+ |
| n00117 | Circular | Flat | Red | S | Entire | Cocci | G+ |
| n00121 | Circular | Raised | Milk White | S | Entire | Cocci | G+ |
| n00123 | Circular | Raised | Milk White | S | Entire | Cocci | G+ |
| n00124 | Circular | Raised | Milk White | S | Entire | Rod-shape | G+ |
| n00125 | Circular | Raised | Yellow | M | Entire | Cocci | G+ |
| n00126 | Circular | Raised | Milk White | S | Entire | Cocci | G+ |
| n00133 | Circular | Raised | Milk White | M | Entire | Cocci | G+ |
| n00134 | Circular | Raised | Milk White | S | Entire | Cocci | G+ |
| n00138 | Circular | Raised | Milk White | S | Entire | Rod-shape | G+ |
| n00139 | Circular | Umbonate | Milk White | M | Entire | Cocci | G+ |
| **JAPONICA endosphere bacteria** | **Form** | **Elevation** | **Color** | **Size** | **Margin** | **Cell morphology** | **G+/G-** |
| n00016 | Circular | Raised | Milk white | M | Entire | Rod-shape | G+ |
| n00017 | Circular | Raised | Milk white | M | Undulate | Cocci | G+ |
| n00018 | Circular | Raised | Milk white | S | Entire | Rod-shape | G- |
| n00019 | Circular | Raised | Milk white | S | Entire | Cocci | G+ |
| n00020 | Circular | Raised | Milk white | S | Undulate | Cocci | G+ |
| n00046 | Circular | Raised | Yellow | L | Entire | Cocci | G+ |
| n00047 | Circular | Umbonate | Milk white | S | Entire | Cocci | G+ |
| n00071 | Circular | Raised | Milk White | M | Entire | Cocci | G- |
| n00072 | Circular | Slightly Raised | Milk White | S | Entire | Cocci | G+ |
| n00073 | Circular | Flat | White | S | Entire | Cocci | G+ |
| n00074 | Circular | Umbonate | White | M | Undulate | Rod-shape | G+ |
| n00075 | Circular | Raised | Milk White | M | Undulate | Cocci | G+ |
| n00076 | Circular | Crateriform | Milk White | M | Entire | Rod-shape | G+ |
| n00077 | Circular | Flat | White | S | Entire | Rod-shape | G+ |
| n00078 | Circular | Flat | White | M | Filiform | Rod-shape | G+ |
| n00079 | Circular | Flat | Milk White | M | Entire | Rod-shape | G+ |
| n00080 | Circular | Flat | White | M | Entire | Rod-shape | G+ |
| n00081 | Circular | Flat | Yellow | S | Entire | Cocci | G+ |
| n00082 | Irregular | Flat | White | L | Undulate | Chain Cocci | G+ |
| n00083 | Irregular | Flat | Cream | L | Filiform | Rod-shape | G+ |
| n00084 | Irregular | Flat | Cream | L | Filiform | Chain Cocci | G+ |
| n00150 | Circular | Umbonate | White | L | Undulate | Chain cocci | G+ |
| n00152 | Circular | Flat | Creame | M | Undulate | Rod-shape | G+ |
| n00153 | Circular | Umbonate | White | S | Entire | Cocci | G+ |
| n00154 | Circular | Slightly raised | White | S | Entire | Cocci | G+ |
| n00155 | Circular | Flat | White | S | Undulate | Rod-shape | G+ |
| n00156 | Circular | Flat | White | S | Entire with transparent margin | Rod-shape | G+ |
| n00164 | Circular | Slightly raised | White | S | Entire | Rod-shape | G+ |
| n00165 | Circular | Raised | Milk White | S | Entire | Cocci | G+ |
| n00174 | Irregular | Slightly Raised | Milk White | S | Lobate | Rod-shape | G+ |
| **JAPONICA phyllosphere bacteria** | **Form** | **Elevation** | **Color** | **Size** | **Margin** | **Cell morphology** | **G+/G-** |
| n00026 | Circular | Flat | Milk white | S | Entire | Cocci | G+ |
| n00027 | Circular | Slightly raised | Milk white | S | Filiform | Rod-shape | G+ |
| n00050 | Circular | Umbonate | Milk white | M | Undulate | Cocci | G- |
| n00051 | Circular | Flat | Milk white | S | Undulate | Cocci | G+ |
| n00052 | Circular | Umbonate | Milk white | M | Entire | Rod-shape | G+ |
| n00053 | Irregular | Slightly Raised | Red | S | Undulate | Rod-shape | G+ |
| n00054 | Circular | Flat | Yellow | S | Entire | Rod-shape | G+ |
| n00055 | Circular | Flat | Milk white | S | Undulate | Rod-shape | G- |
| n00088 | Circular | Flat | Milk White | M | Entire with transparent margins | Cocci | G+ |
| n00089 | Circular | Flat | Milk White | M | Entire with transparent margins | Chain Cocci | G+ |
| n00162 | Circular | Raised | Yellow | S | Entire | Diplo cocci | G+ |
| n00169 | Circular | Raised | White | S | Entire | Rod-shape | G+ |
| n00170 | Circular | Raised | Milk White | S | Entire | Rod-shape | G- |
| n00170-1 | Irregular | Raised | Milk White | M | Undulate | Rod-shape | G+ |
| n00170-2 | Circular | Raised | Milk White | M | Entire | Rod-shape | G+ |
| n00171 | Circular | Raised | Yellow | S | Entire | Diplo cocci | G+ |
| n00172 | Circular | Flat | White | S | Entire | Rod-shape | G+ |
| n00175 | Circular | Raised | Yellow | S | Entire | Diplo cocci | G+ |

**Table S5.** Phosphate solubilizing bacteria (PSB) with ACC deaminase production. “Strong +” indicates PSB grew strongly on DF minimal salt medium supplemented with 3 mM ACC as sole nitrogen source, while “Slight +” indicates weak growth. “None” means no growth.

| PSB codes | ACC deaminase (overall 3 trials) |
| --- | --- |
| n00007 | Slight + |
| n00008 | None |
| n00009 | Slight + |
| n00010 | Strong + |
| n00011 | Strong + |
| n00014-1 | Strong + |
| n00014-2 | Strong + |
| n00015 | Strong + |
| n00016 | Strong + |
| n00024-1 | Strong + |
| n00024-2 | Strong + |
| n00064 | Slight + |
| n00069 | Slight + |
| n00078 | None |
| n00080 | None |
| n00082 | Slight + |
| n00086 | Slight + |
| n00093 | None |
| n00128 | Strong + |
| n00132 | Slight + |
| n00170 | Slight + |
| n00170-2 | Strong + |
| n00172 | Slight + |
| n00163 | Slight + |
| n00167 | Slight + |
| *E. coli* strain OP50 | None |
| *Agrobacterium* | Strong + |
| *Serratia marcescens* | Slight + |

**Table S6.**
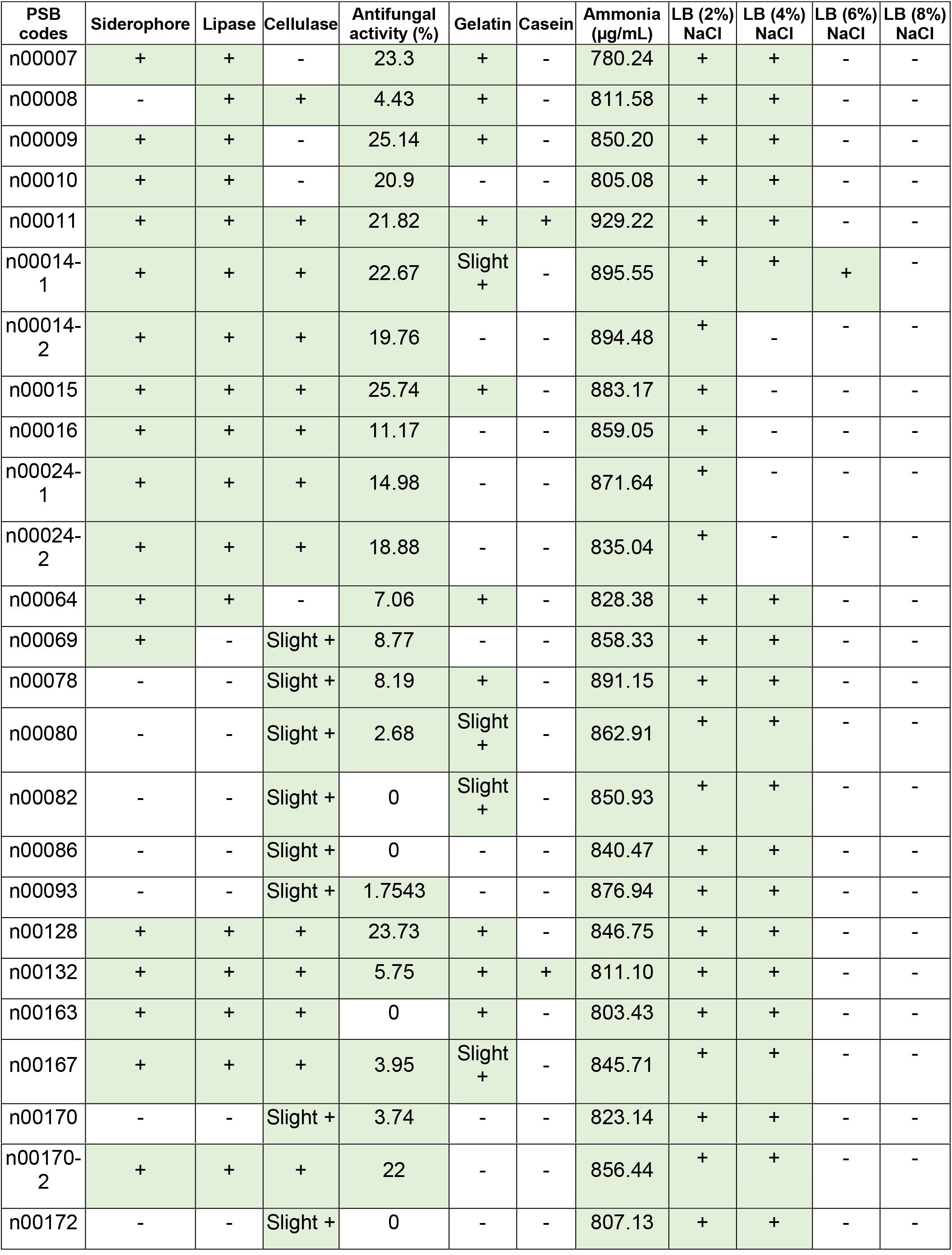
Indirect plant growth promoting assays and *in vitro* assay results for phosphate solubilizing bacteria (PSB) salt tolerance.

**Table S7.** Bacterial phenotypic fingerprint of carbon source usage (Biolog microplates). “+” indicates bacterial use of the specific carbon source and “-” indicates that bacteria did not grow in the microplate. n00132, *Pseudomonas mosselii;* n00163, *Paenibacillus rigui*; n00167, *Brevibacillus sp900114075*; n00170, *Microvirga sp003151255*; n00172, *Paenibacillus graminis*.

| Carbon Sources | Agro | n00132 | n00163 | n00167 | n00170 | n00172 |
| --- | --- | --- | --- | --- | --- | --- |
| Dextrin | + | + | + | + | + | + |
| D-Maltose | + | + | + | + | + | + |
| D-Trehalose | + | + | + | + | + | + |
| D-Cellobiose | + | + | + | + | + | + |
| Gentiobiose | + | + | + | + | + | + |
| Sucrose | + | + | + | + | + | + |
| D-Turanose | + | + | + | + | + | + |
| Stachyose | + | - | - | - | - | + |
| D-Raffinose | + | - | - | - | - | - |
| α-D-Lactose | + | - | - | - | + | + |
| D-Melibiose | + | - | - | - | - | + |
| β-Methyl-D-Glucoside | + | + | + | + | + | + |
| D-Salicin | + | + | + | + | + | + |
| N-Acetyl-D-Glucosamine | + | + | + | + | + | + |
| N-Acetyl-β-D-Mannosamine | + | + | + | + | + | + |
| α-D-Glucose | + | + | + | + | + | + |
| D-Mannose | + | + | + | + | + | + |
| D-Fructose | + | + | + | + | + | + |
| D-Galactose | + | + | + | + | + | + |
| D-Fucose | + | - | - | - | - | - |
| L-Fucose | + | - | - | - | - | - |
| L-Rhamnose | + | - | - | - | - | - |
| Inosine | + | + | + | + | + | + |
| D-Sorbitol | + |  |  |  |  |  |
| D-Mannitol | + | + | + | + | + | + |
| D-Arabitol | + | - | - | - | - | - |
| myo-Inositol | + | - | - | - | - | - |
| Glycerol | + | - | + | + | + | + |
| D-Glucose-6-PO<sub>4</sub> | + | - | - | - | - | - |
| **D-Fructose-6-PO<sub>4</sub>** | + | - | - | - | - | - |
| **D-Aspartic Acid** | + | - | - | - | - | - |
| **L-Alanine** | + | - | - | - | - | - |
| **L-Aspartic Acid** | + | - | - | - | - | - |
| **L-Glutamic Acid** | + | - | - | - | - | - |
| **L-Pyroglutamic Acid** | + | - | - | - | - | - |
| **L-Serine** | + | - | - | - | - | - |
| **Pectin** | + | + | + | + | + | + |
| **D-Galacturonic Acid** | + | - | - | - | - | - |
| **L-Galactonic Acid Lactone** | + | - | - | - | - | - |
| **D-Gluconic Acid** | + | + | + | + | + | + |
| **D-Glucuronic Acid** | + | - | - | - | - | - |
| **Glucuronamide** | + | - | - | - | - | - |
| **Mucic Acid** | + | - | - | - | - | - |
| **Quinic Acid** | + | - | - | - | - | - |
| **Methyl Pyruvate** | + | - | + | + | + | - |
| **D-Lactic Acid Methyl Ester** | + | - | - | - | - | - |
| **L-Lactic Acid** | + | + | + | + | + | + |
| **D-Malic Acid** | + | - | - | - | - | - |
| **L-Malic Acid** | + | - | - | - | - | - |
| **Bromo-Succinic Acid** | + | - | - | - | - | - |
| **Tween 40** | + | + | + | + | + | + |
| **Acetoacetic Acid** | + | + | + | + | + |  |
| **Propionic Acid** | + | - | - | - | - | - |
| **Acetic Acid** | + | - | - | - | - | - |

**Table S8.**
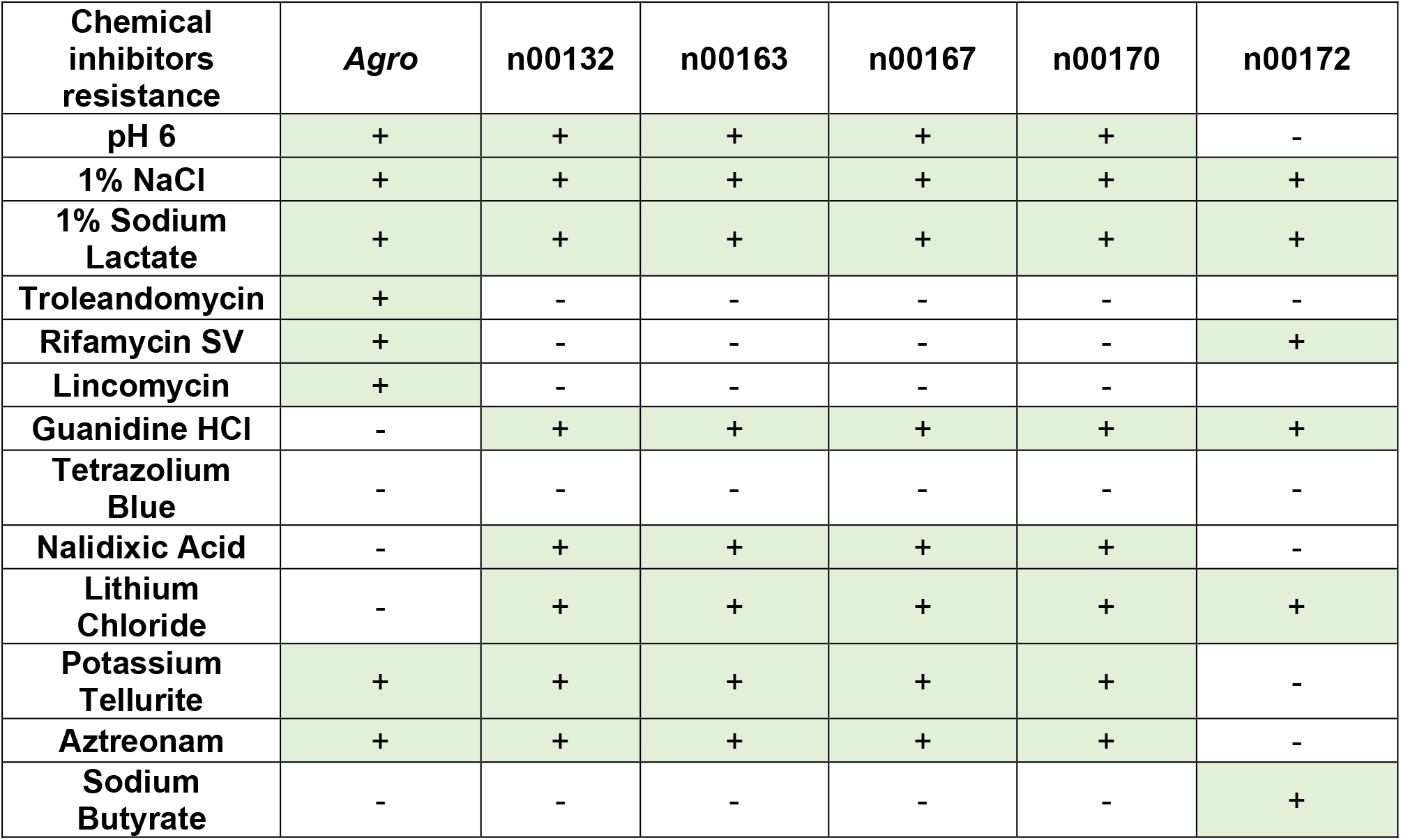
Bacterial phenotypic fingerprint of chemical sensitivity (Biolog microplates). “+” indicates bacterial resistance to a specific chemical and “-” indicates bacterial sensitivity to a specific chemical.

**Table S9.** Growth inhibition by n00172 (*Paenibacillus graminis*). Two antibacterial assays, overlay-agar and cross streak, show n00172’s ability to inhibit the growth of the four other phosphate solubilizing bacteria (PSB). In overlay-agar test, “+” indicates presence of an inhibition halo around n00172 when each of these four PSB are overlaid. In the cross streak test, growth inhibition of the four other PSB when crossing the n00172 bacterium is shown in mm (the test was done twice). n00132, *Pseudomonas mosselii*; n00163, *Paenibacillus rigui*; n00167, *Brevibacillus sp900114075*; n00170, *Microvirga sp003151255*.

| Antibacterial activity/Bacteria | n00132 | n00163 | n00167 | n00170 | n00172 |
| --- | --- | --- | --- | --- | --- |
| Overlay-agar | - | - | - | - | + |
| Cross-streak | 2 mm/1 mm | 3 mm/1 mm | 1 mm/1 mm | 1 mm/1 mm | - |

**Table S10.** Secondary metabolite regions of isolate n00172 (*Paenibacillus graminins*) identified using strictness ‘relaxed’. NRPS = Non-ribosomal peptide synthetase cluster, Type I PKS = Type I Polyketide synthase.

| Region | Type | From | To | Most similar known cluster | Similarity |
| --- | --- | --- | --- | --- | --- |
| 3.1 | NPRS, T1PKS, NRPS-like | 181,635 | 238,230 |  |  |
| 4.1 | cyclic-lactone-autoinducer | 812 | 21,351 |  |  |
| 4.2 | RRE-containing | 55,265 | 76,665 |  |  |
| 5.1 | cyclic-lactone-autoinducer | 34,965 | 55,683 |  |  |
| 7.1 | NPRS, T1PKS | 1 | 38,457 | xenocoumacin 1 / xenocoumacin II, NRP + Polyketide:Modular type I | 28% |
| 10.1 | NPRS, transAT-PKS | 154,535 | 193,126 |  |  |
| 13.1 | cyclic-lactone-autoinducer | 150,188 | 161,901 |  |  |
| 20.1 | NPRS, transAT-PKS, T1PKS, NRPS-like | 1 | 97,214 |  |  |
| 30.1 | lassopeptide | 17,331 | 41,158 | Paeninodin, RiPP | 100% |
| 33.1 | RRE-containing | 25,743 | 47,170 |  |  |
| 51.1 | cyclic-lactone-autoinducer | 4,648 | 25,296 |  |  |
| 55.1 | cyclic-lactone-autoinducer | 16,231 | 36,755 |  |  |
| 69.1 | NRPS-like | 1 | 13,082 |  |  |
| 71.1 | NPRS | 1 | 10,121 |  |  |
| 76.1 | transAT-PKS-like | 1 | 4,361 |  |  |
| 77.1 | NPRS | 1 | 2,877 |  |  |
| 83.1 | NPRS | 1 | 1,397 |  |  |

